# Spinal inhibitory neurons degenerate before motor neurons and excitatory neurons in a mouse model of ALS

**DOI:** 10.1101/2023.09.17.558103

**Authors:** Roser Montañana-Rosell, Raghavendra Selvan, Pablo Hernández-Varas, Jan M. Kaminski, Dana B. Ahlmark, Ole Kiehn, Ilary Allodi

## Abstract

Amyotrophic lateral sclerosis (ALS) is characterized by the progressive loss of somatic motor neurons. Major focus has been directed to motor neuron intrinsic properties as a cause for degeneration while less attention has been given to the contribution of spinal interneurons. In the present work, we applied multiplexing detection of transcripts and machine learning-based image analysis to investigate the fate of multiple spinal interneuron populations during ALS progression in the SOD1^G93A^ mouse model. The analysis showed that spinal inhibitory interneurons are affected early in disease, before motor neurons death, and are characterized by a slow progressive degeneration, while excitatory interneurons are affected later with a steep progression. Moreover, we report differential vulnerability within inhibitory and excitatory subpopulations, with interneurons directly projecting onto motor neurons being preferentially affected. Our study reveals a strong interneuron involvement in ALS development with interneuron specific degeneration. This points to differential involvement of diverse spinal neuronal circuits that eventually may be determining for motor neuron degeneration.

**Teaser:** A new approach to study the motor neuron disorder Amyotrophic Lateral Sclerosis shows the early and differential dysregulation of spinal interneurons.

## Introduction

In Amyotrophic lateral sclerosis (ALS) somatic motor neurons degenerate leading to muscle denervation and wasting (1). Hence, subjects affected by the disease progressively lose the ability to control muscles and perform movements. Execution of movements requires synchronized or reciprocal activation of motor neuron pools for appropriate muscle activation during intra- and interlimb coordination, flexor-extensor and left-right alternations (2). Such activation is regulated by a complex network of inhibitory and excitatory intraspinal neurons decoding the information received from the descending tracts and the afferent pathways (3). Within the ventral horn of the spinal cord these inhibitory and excitatory neurons can be divided into four cardinal classes: V0, V1, V2 and V3, identified by the expression of specific molecular markers defining cell fate during development (4). Knowledge generated by research in the motor control field has shown that, in vertebrates, specific subpopulations of inhibitory (expressing glycine and GABA) and excitatory (expressing glutamate and acetylcholine) spinal neurons form the central pattern generators (CPGs), controlling rhythmic movements, and regulating different aspects of locomotion (2, 3, 5, 6).

These cardinal classes are further diversified. Thus, V0 interneurons can be further divided based on their axonal projections, and excitatory or inhibitory neurotransmitters. The V0_C/G_ subpopulations are ipsilaterlly projecting and contain either acetylcholine or glutamate, respectively. Found around the central canal, they are characterized by expression of the Pitx2 marker, and control changes in muscle tone (7). The largest population of V0 interneurons are contralaterally projecting either being excitatory (V0_V_) or inhibitory (V0_D_) and characterized by expression of Dbx1 and Evx1 transcription factors. They control the left-right coordination at slow and high speed of locomotion, respectively (8-10). V1 interneurons are a large ipsilaterally projecting inhibitory population. In the adult mouse 80% of them are glycinergic, although often co-expressing GABA (11). Previous studies have shown that this population is highly heterogeneous and, depending on marker co-expression, up to 50 subpopulations can be identified (12, 13). In mammals, V1 interneurons are known to control the speed of locomotion (14), and Renshaw cells as well as Ia interneurons belong to the V1 class (11). V2 interneurons are also ipsilaterally projecting and can be divided in V2a and V2b, V2a being excitatory and V2b inhibitory neurons. V2a neurons are characterized by expression of Chx10 transcription factor (15), and are rhythmically active during locomotion (16). V2b neurons are identified by Gata2/3 expression (17), and together with V1 interneurons are required for coordination of flexor-extensor alternation (18, 19). The V3 interneurons projects both contralaterally and ipsilaterally (20, 21) and are excitatory; ablation or silencing of those neurons leads to changes in left-right coordination (20). Two other molecularly-defined excitatory populations are the Shox2 (16) and the Hb9 (also called Mnx1) (22) positive neurons. Shox2 neurons are found in the intermediate area of the spinal cord, 75% of them co-expressed Chx10, while the remaining 25% which are Shox2 positive and Chx10 negative are known to generate locomotor activity (16). Hb9 is expressed by motor neurons as well as many interneurons in the intermediate are and more ventral which also contribute to locomotor rhythm generation (22). Growing evidence has reported spinal circuit dysregulations in ALS (for review (23)). The V0-cholinergic Pitx2+ (V0_C_) neurons are the main source of C boutons, which are cholinergic synapses found on motor neurons and described to undergo changes during ALS progression (24). Electrophysiological changes have been reported in glycinergic interneurons already at early postnatal stages, with the most ventrally located neurons being less excitable (25). Our recent findings showed that already at postnatal day 45, there is a loss of glycinergic synapses on fast motor neurons (26). Further analysis revealed that V1 interneurons are affected in the SOD1^G93A^ mice (27) before motor neuron degeneration and muscle denervation (26). Other studies performed on SOD1 mouse models as well, reported degeneration of V2a neurons at late disease stages (28, 29). However, the exact temporal dynamics of degeneration of the essential spinal motor related circuits throughout disease development has not been revealed. Since the interneuron populations are essential for all movement we set out to perform such studies.

A clear challenge to this type of investigation is the inherent difficulties in being able to study the fate of multiple interneuron subpopulations during disease progression. To succed it requires the simultaneous visualization of several molecular markers in in the spinal cord as the disease progress. Previous studies have utilized multiplexed antibody staining with up to four different markers to identify neuronal subpopulations within the spinal cord (12). However, recent advancement in the multiplexing detection of transcripts now allow for the visualization of tens and up to hundreds of markers at the same time (30, 31). While multiplexing detection and visualization is now routinely performed in several laboratories, reliable quantification of multiple transcripts in the same cells has been proven difficult and often requires manual annotation. Hence, in the present study we developed machine learning-based tools to automize cell segmentation and subsequent transcript registration, as well as a bioinformatic analysis pipeline to perform spatial analysis of differentially expressed transcripts. The combination of the multiplexing detection and the computational analysis allowed us to investigate the changes in expression of molecular markers identifying multiple classes of interneurons in the spinal cord as well as motor neuron populations in the SOD1^G93A^ mouse at three time points of disease progression: pre-symptomatic – postnatal day (P) 30, onset of locomotor phenotype – P63, and symptomatic with motor neuron death – P112. We find that inhibitory subpopulations of interneurons start losing their specific markers (e.g., Engrailed-1 and Calb1) early in disease. In contrast, excitatory interneurons (e.g., Chx10 positive) lose their specific markers at late stages in disease, although these changes in expression appear quickly at symptomatic stages. Interestingly, interneurons directly projecting on to motor neurons seem to be affected earlier in the SOD1^G93A^ mice. Taken together, these results point towards a larger contribution of spinal circuits in disease, with distinct degenerative temporal dynamics between inhibitory and excitatory interneurons.

## Results

### Developmentally defining neuronal markers are maintained within the adult spinal cord and can be used to identify interneurons in health and disease

The expression of molecularly defining interneuron markers has been mainly characterized during mouse development at embryonic and postnatal stages (4, 12). Among these markers are transcription factors, whose expression appear to be downregulated in the adult tissue making antibody detection less sensitive. We previously showed that the Engrailed-1 transcript, identifying V1 interneurons, can be reliably detected in the mouse adult spinal cord by *in situ* hybridization (26, 32). In the present study, we compared expression of 24 markers (Fig. 1A-B) in P1 and P28 mouse spinal cord tissue to validate their presence and spatial location in the adult, by utilizing a *in situ* sequencing technique (CARTANA), now replaced by Xenium (10X Genomics)(Fig. 2A) (31). This powerful technique allows for the detection of more than 250 markers at the same time, by utilizing barcoded probes detecting the specific transcripts (31). Upon sequencing, information about the spatial localization of the barcodes is provided as X-Y coordinates within the acquired tile images (31) which allow for exact localization of transcript within the spinal cord (Fig. 2A). However, for exact subpopulation identification, co-expression of multiple markers is often required. Therefore, outline of neuronal soma or cell bodies are needed in order to be able to assign potential expression of multiple markers within the same cell. Previous computational methods have been developed to identify cell body boundaries based on segmentation of nuclear (DAPI) staining (33, 34). However, neurons found within the ventral and dorsal areas of the spinal cord differ significantly in shape and size, with ventral neurons being larger and dorsal neurons being characterized by smaller and rounder somas. Therefore, segmentation based on nuclear stain might not capture the diverse cell body size in the spinal cord. To segment spinal neurons, we therefore implemented a machine learning deep ensemble approach as shown in Fig. 2A. To get an outline of cell bodies we used NeuroTrace, which provides a fluorescent Nissl staining of the cytoplasm of neurons and nuclei. NeuroTrace images of the analyzed spinal cord tissue were obtained after *in situ* sequencing and registered to the DAPI images obtained by CARTANA for X-Y coordinate registration (Fig. 2A). Probabilistic segmentation of the cell bodies was then obtained based on the NeuroTrace staining, utilizing an updated version of a deep ensemble method previously developed for limited labelled training data (35). Neuron annotations for both P1 (n = 52 annotated patches, from N = 2 sections from 2 independent mice) and P28 (n = 20 annotated patches, from N = 2 sections from 2 independent mice) tissue were generated and used to train parallel models using U-Net Convolutional Neural Networks (CNN) with k-fold cross-validation (Fig. 2A). The final ensemble model obtained from the combination of P1 and P28 models was used for probabilistic segmentation of the tile images. A threshold of 0.4 confidence and a structural kernel of 5 was applied to define the final segmentation used for analysis (neurons visualized in yellow and green). Transcript X-Y coordinates were then registered to segmented neurons, and spatial analysis was performed per hemicord. A total of N = 6 P1 and N = 29 P28 spinal cord sections (from 3 and 11 mice, respectively) was analyzed to validate transcript expression and spatial localization. Cross sections obtained from lumbar (L) segments 1-3 were included in the study. Fig. 2B shows an example of spatial analysis for the cardinal neuronal identity markers Pitx2, En1, Chx10, Shox2, and choline acetyltransferase (ChAT) in P1 vs P28 tissue (the graph includes pooled data from 2 spinal cords from N= 1 mice at P1 and N= 2 mice at P28). An example of spatial localization of putative markers for the V1 subpopulations including Calbindin (Calb1), Calretinin (Calb2) (11), Foxp2, Pou6f2, and Sp8 (12) is shown in Fig. 2C. Examples of combinatorial expression of interneuron markers are shown in Fig. S1A-B.

**Fig. 1.**
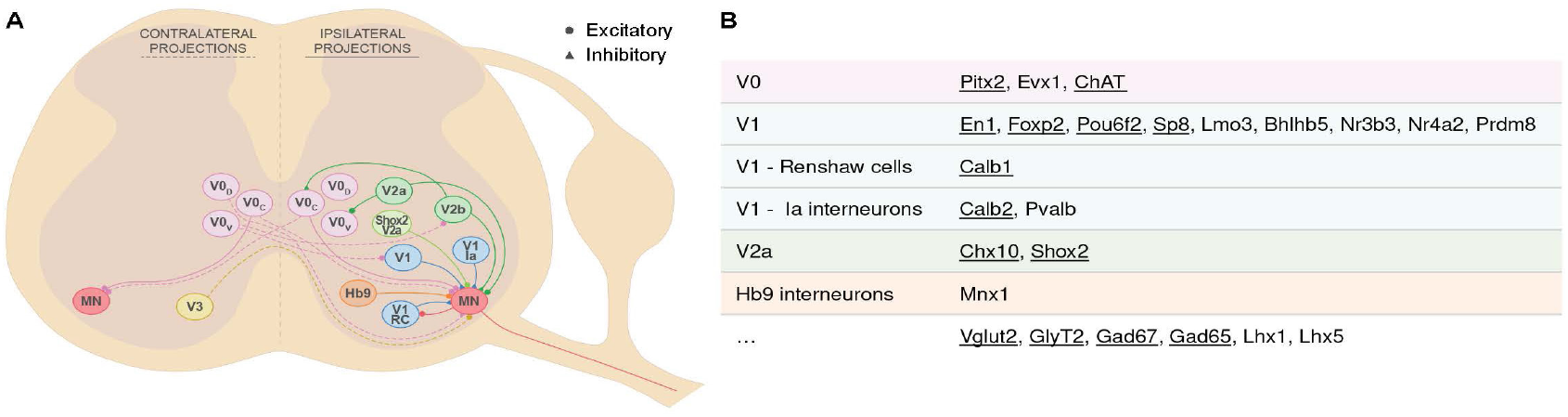
Interneuronal markers investigated in the present study and their circuitry. (A) Schematic of the locomotor circuits in the mammalian lumbar spinal cord. Shown are the cardinal ventral interneuron subpopulations (V0_D_, V0_V_, V0_C_, V1, V1 Renshaw cells (RC), V1 Ia interneurons, V2a, V2b, V3) as well as Shox2 and Hb9 interneurons, and their main ipsilateral (solid) or contralateral (dashed) projections. Inhibitory inputs are shown as triangles, excitatory as circles. (B) Panel of markers detected throughout the study representing the different interneuron populations of interest. Transcript expression of underlined markers was quantified for characterization of interneuron dysregulation in ALS progression.

**Fig. 2.**
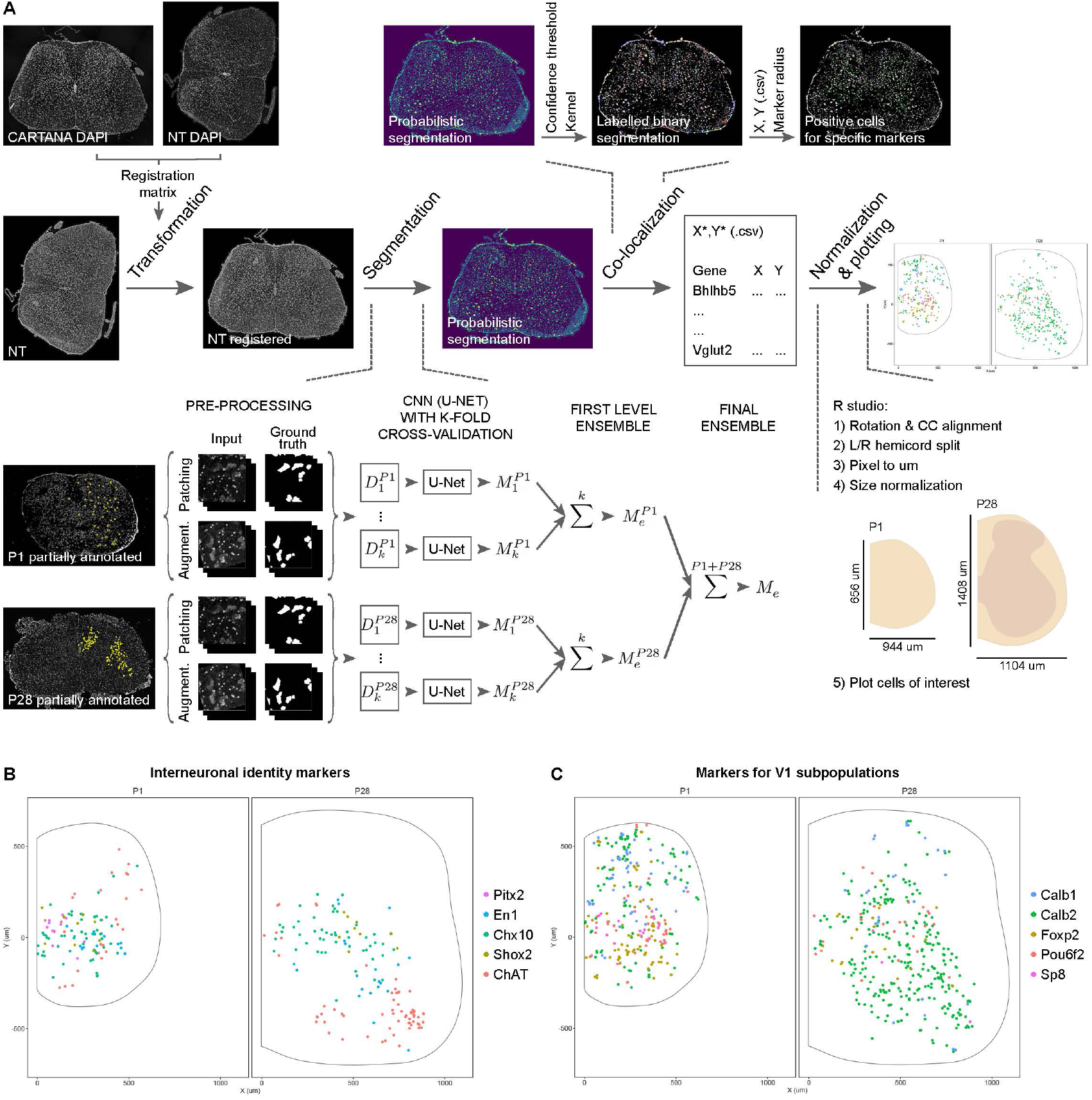
Fate defining markers are present in adult mouse tissue. (A) Pipeline used for processing and analysis of in situ sequencing data. NeuroTrace (NT) images were acquired and registered to in situ sequencing data based on DAPI staining. These were then used for segmentation of neuronal cells using a machine learning deep ensamble model based on the use of U-Net convolutional neural network with k-fold cross-validation, which were applied separately to patches (512×512 px) of annotated data from P1 (52 annotated patches) and P28 (20 annotated patches) and merged through two ensamble steps to obtain probabilistic segmentation. A confidence threshold and structural kernel were applied to obtain final segmentation, to which in situ sequencing coordinates were registered. Only transcripts that localized to segmented cells were used for final processing in R, where coordinates were normalized and plotted for visualization and qualitative analysis. (B) Validation of identity markers by in situ sequencing technique in wild-type young adult mouse lumbar spinal cord (P28, right) compared to early postnatal (P1, left), for the interneuron populations included in the study: Pitx2 (pink) for V0_C/G_, En1 (blue) for V1, Chx10 (green) for V2a, Shox2 (khaki), and ChAT (salmon) for V0_C_ and motor neurons. (C) Validation of markers in adult mouse tissue for the V1 interneuron subpopulations including Calb1 (blue) putative for Renshaw cells, Calb2 (green) putative for Ia, and the cardinal clade markers Foxp2 (khaki), Pou6f2 (salmon), and Sp8 (pink). Data in (D-E) are pooled from n = 2 sections from N = 1 mice for P1, and N = 2 mice for P28.

The analysis demonstrates that the interneuron markers that have been used embryonically and early postnatally to characterize spinal interneuron populations (4, 36, 37) are also found in the adult spinal cord. Based on the spatial localization it appears that the marker-identified interneuron populations maintain their identity in the adult spinal cord.

### Investigation of inhibitory and excitatory interneuron markers in the SOD1G93A mouse model reveals larger contribution of neural circuits in disease

For single mRNA molecule quantifications in healthy and ALS tissue, the RNAscope HiPlexUp technique (ACD, BioTechne) was utilized (Fig. 3, Fig. S2A). RNAscope is known to detect ∽95% of mRNA molecules of the targeted transcripts, and the HiPlexUp technique allows for simultaneous detection of up to 48 transcripts in the same tissue. Probes for the 24 transcripts investigated by *in situ* sequencing (Fig. 1B) were designed and validated by ACD BioTechne. Procedure was performed as shown in Fig. S2A. Upon two series of Hybridization-Amplification and four labeling and cleaving rounds per series, images were acquired in a total of 8 rounds (3 probes imaged per round). Also in this case, NeuroTrace and DAPI staining were included for image registration and cell segmentation. Using ImageJ, registration matrixes for the 8 HiPlexUp rounds plus NeuroTrace were generated based on DAPI staining, and these were used to transform and merge 26 superimposed tile images of each spinal cord section (one per transcript + DAPI + Neuro Trace) which were used for further processing in the ZEN software (Zeiss). Here, the machine learning based Intellesis tool (Zeiss) was utilized for cell segmentation and neuron enrichment (Fig. S2B-C, in orange is the positive fraction = neurons included in the study, in blue is the negative fraction = background and other cells). X-Y coordinates and area were obtained for each segmented neuron, while the intensity mean and standard deviation were obtained for each transcript expressed within each neuron (Fig. S2B).

**Fig. 3.**
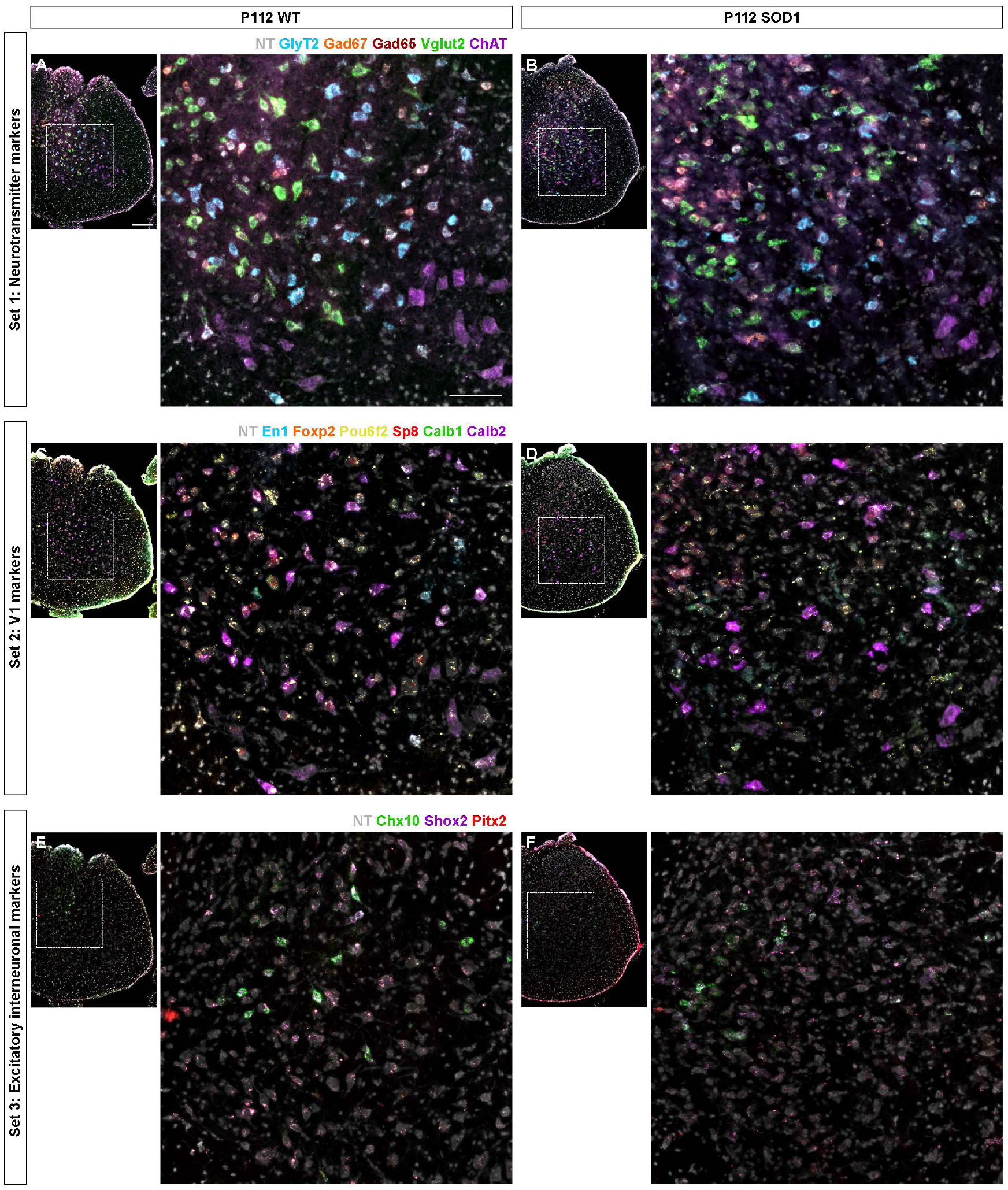
Interneuron marker transcript detected by RNAscope HiPlexUp. Microscopy images obtained by RNAscope HiPlexUp multiplexed in situ hybridization in wild-type (wt) (left) and SOD1^G93A^ mice (right) at P112, including all markers used for characterization of interneuron dysregulation. To facilitate visualization, markers have been divided in three subsets. For each subset and condition, an overview of a full hemisection is shown, together with an inset with higher magnification. (A-B) Subset 1 includes neurotransmitter markers: GlyT2 (blue), Gad67 (orange), Gad65 (deep red), Vglut2 (green), and ChAT (purple). (C-D) Subset 2 includes markers for the V1 population: En1 (blue), Foxp2 (orange), Pou6f2 (yellow), Sp8 (red), Calb1 (green), and Calb2 (purple). (E-F) Subset 3 includes markers for some excitatory populations: Chx10 (green, for V2a), Shox2 (purple), and Pitx2 (red, for V0_C/G_). Scale bars, 200 μm for hemisections, 100 μm for insets. NT counterstaining in grey.

Three different timepoints were included in the study, P30 (pre-symptomatic), P63 (onset of locomotor phenotype) and P112 (symptomatic with motor neuron death); 3 wildtype (wt) and 3 SOD1^G93A^ mice were included per timepoint. Analysis was performed per hemicord, since SOD1^G93A^ show asymmetrical degeneration within the spinal cord (similarly to ALS patients with spinal onset (38)). Both left and right hemicords were analyzed for each mouse, with 2-5 hemisections included per hemicord. With this approach, 98.943 neurons were segmented, of which 84.791 were included in the study after applying a size threshold of 70 μm2. When comparing the number of segmented neurons per condition and timepoint, we observed that comparable number of neurons were included in the study, with an average of 625 segmented cells per hemisection (multiple unpaired t tests with Welch correction; P30: t = 0.5782, df = 38, P > 0.9999; P63: t = 1.311, df = 41, P = 0.5918; P112: t = 0.09803, df = 46, P > 0.9999; n = 20-26 hemisections, N = 3 mice per condition) (Fig. S2D, the spatial distribution of the segmented neurons is shown for individual mice in Fig. S2E). Interestingly, at P112, changes in expression levels are visually notable for some of the markers (e.g., En1 and Chx10), due to the clear decrease in intensity when compared to control conditions (Fig. 3C-D and E-F). All 24 markers were investigated in disease progression in the SOD1^G93A^ mouse. However, in the present work, only the following markers related to major classes of interneuorns will be discussed (underlined in Fig. 1B): En1, FoxP2, Pou6f2, Sp8 for V1 interneurons; Calb1 for V1 putative Renshaw Cells; Calb2 for V1 putative Ia interneurons; Chx10, Shox2 for V2a interneurons; Pitx2, ChAT, Vglut2 for V0_C/G_ interneurons; ChAT for motor neurons; GlyT2, Gad67, Gad65 and vGlut2 for general neurotransmitters.

### V1 inhibitory interneurons are affected early in disease and show differential vulnerability among subpopulations

Upon image postprocessing (Fig. 4A, Fig. S2A-B), V1 interneurons were defined as inhibitory neurons (GlyT2, Gad65 and/or Gad67 positive), positive for En1 (Fig. 4B-C). Analysis performed at pre-symptomatic stage did not show alterations in En1 expression (P30: nested unpaired two-tailed t test, t=1.305, df=10, P=0.2210, n=20-22 hemisections and N=6 hemicords from 3 mice per condition) (Fig. 4D). Our previous data with manual detection showed ∽25% decrease in the En1 transcript expression already at P63 in the SOD1^G93A^ mice (26). In the present study, utilizing an automized approach, we were able to validate similar decrease in expression at P63 (P63: 19.2% downregulation, nested unpaired two-tailed t test, t= 2.493, df= 10, P= 0.0318, n= 21-22 hemisections and N= 6 hemicords from 3 mice per condition) (Fig. 4D). The En1 loss was further intensified in symptomatic mice with a total decrease of 45.4% compared to controls (P112: nested unpaired two-tailed t test, t= 6.379, df= 47, P < 0.0001, n= 23-26 hemisections and N= 6 hemicords from 3 mice per condition), which was also evident upon visualization of spatial distribution and count-based kernel density estimation (Fig. 4B-E).

**Fig. 4.**
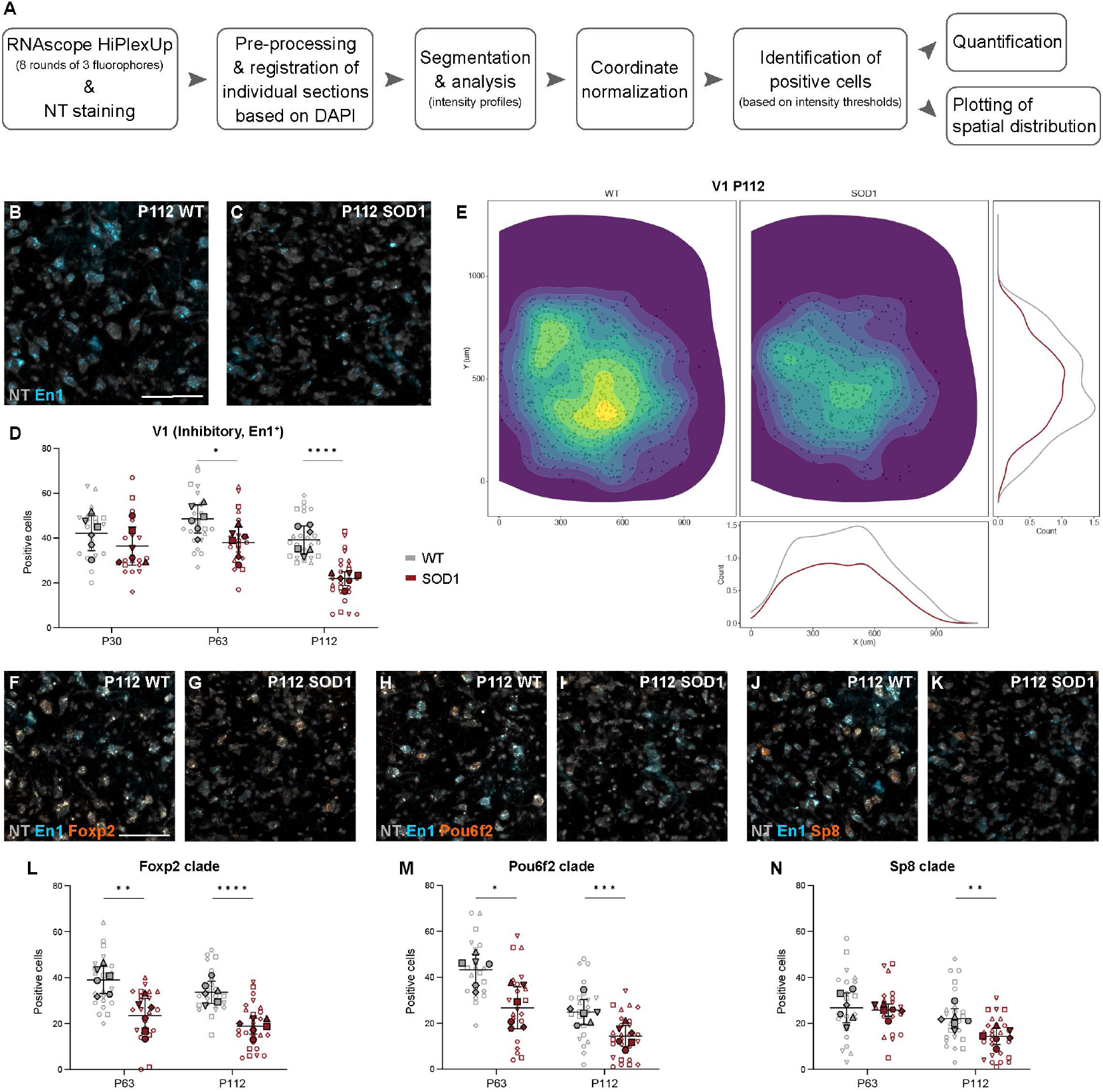
The V1 interneuron population and its cardinal clades are dysregulated in the SOD1^G93A^ mouse. (A) Transcript detection, image processing and cell analysis workflow. For further details check Fig. S2. (B-C) Microscopy images from RNAscope HiPlexUp in situ hybridization assay showing detection of En1 transcript (blue) with NeuroTrace counterstaining (NT, grey) in wild-type (wt) (B) and SOD1^G93A^ (C) lumbar spinal cord at P112. (D) Significant reduction in number of inhibitory (GlyT2+, Gad67+ and/or Gad65+) En1+ neurons in SOD1^G93A^ mice (red) compared to healthy control mice (grey) at P63 and P112 (nested unpaired two-tailed t tests; P30 P = 0.2210, P63 P = 0.0318, P112 P < 0.0001). (E) Spatial distribution within the spinal cord at P112 shows the loss of inhibitory En1+ neurons in the SOD1^G93A^ mouse (right/red) compared to healthy control mice (left/grey), which is especially evident in the ventral-most region. Data were pooled from all P112 sections and shown as individual cells (black dots) with count-based kernel density estimations in 2E (main panel), and in X (bottom panel) and Y dimensions (right panel). (F-K) RNAscope HiPlexUp microscopy images of En1 transcript (blue) co-localized with V1 cardinal clade markers (orange) Foxp2 (E-F), Pou6f2 (G-H) and Sp8 (I-J) with NT (grey), in healthy control mice and SOD1^G93A^ mice at P112. (L-N) Quantification in SOD1^G93A^ mice versus healthy control mice shows significant decrease in Foxp2+ neurons at P63 and P112 (L) (nested unpaired two-tailed t tests; P63 P = 0.0036, P112 P < 0.0001), Pou6f2+ neurons at P63 and P112 (M) (nested unpaired two-tailed t tests; P63 P = 0.0100, P112 P = 0.0007), and Sp8+ neurons at P112 only (N) (nested unpaired two-tailed t tests; P63 P = 0.7444, P112 P = 0.0083). Scale bars, 100 μm. 2D kernel densities in (E) were plotted with 10 bins using viridis scale. N = 6 hemicords from 3 mice (filled shapes). Number of hemisections (empty shapes): P30 n(wt) = 22, n(SOD1) = 20; P63 n(wt) = 22, n(SOD1) = 21; P112 n(wt) = 23, n(SOD1) = 26. Data shown as mean ± SD.

We then investigated potential differential vulnerability within V1 subpopulations by analyzing three main clades defined by the expression of FoxP2, Pou6f2 and Sp8 (12) at P63 and P112 timepoints (Fig. 4F-N). Quantified neurons were positive for GlyT2, Gad65, Gad67 and En1 markers as well as FoxP2, Pou6f2 or Sp8 (Fig. 4F-K). The analysis revealed a differential vulnerability among clades, with FoxP2 (P63: 37.4% downregulation, nested unpaired two-tailed t test, t= 3.777, df= 10, P= 0.0036, n= 22-21 hemisections and N= 6 hemicords from 3 mice per condition) (Fig. 4L) and Pou6f2 (P63: 35.1% downregulation, nested unpaired two-tailed t test, t= 3.168, df= 10, P= 0.0100, n= 22-21 hemisections and N= 6 hemicords from 3 mice per condition) (Fig. 4M) positive neurons decreasing in number earlier compared to the Sp8 positive ones (P63: ns, nested unpaired two-tailed t test, t= 0.328, df= 41, P= 0.7444, n= 22-21 hemisections and N= 6 hemicords from 3 mice per condition) (Fig. 4N). Despite appearing initially more resistant, Sp8 positive neurons are reduced in numbers at P112 (P112: 36.4% downregulation, nested unpaired two-tailed t test, t= 2.754, df= 47, P= 0.0083, n= 23-26 hemisections and N= 6 hemicords from 3 mice per condition) (Fig. 4N). At the same timepoint, FoxP2 and Pou6f2 are further decreased in numbers (P112 FoxP2: 44.3% downregulation, nested unpaired two-tailed t test, t= 5.502, df= 47, P < 0.0001; P112 Pou6f2: 43.9% dowregulation, nested unpaired two-tailed t test, t= 3.637, df= 47, P= 0.0007; n= 23-26 hemisections and N= 6 hemicords from 3 mice per condition) (Fig. 4L-M). Spatial information regarding changes of expression at P63 of En1 transcripts and the three clades is shown in Fig. S3A-D.

Renshaw cells and Ia interneurons, members of the V1 subpopulations, were also analysed at P63 and P112. These functionally defined populations are found in specific areas of the ventral spinal cord (11). In this study, putative Ia interneurons were identified as inhibitory interneurons (GlyT2+, Gad65+ and/or Gad67+), positive for En1 and Calb2 (also known as Calretinin) (11) (Fig. 5A-B). Moreover, their location was restricted to the ventro-lateral area of the spinal cord, where Ia interneurons are found (Fig. 5D). Decrease in inhibitory En1+/Calb2+ neuron numbers was found at P63 (28.0% downregulation) as well as P112 (60.4% downregulation) (Fig. 5C-D) (P63: nested unpaired two-tailed t test, t= 2.022, df= 41, P= 0.0498, n= 22-21 hemisections and N= 6 hemicords from 3 mice per condition; P112: nested unpaired two-tailed t test, t= 4.291, df= 10, P= 0.0016, n= 23-26 hemisections and N= 6 hemicords from 3 mice per condition). Putative Renshaw cells, instead, were identified by expression of the inhibitory markers GlyT2, Gad65 and/or Gad67, in combination with En1 and Calb1 (Fig. 5E-F) and their ventro-medial location in the spinal cord (Fig. 5H). The number of ventral inhibitory En1+/Calb1+ neurons were found to be 52.8% reduced at P63 (P63: nested unpaired two-tailed t test, t= 2.587, df= 10, P= 0.0271, n= 22-21 hemisections and N= 6 hemicords from 3 mice per condition) (Fig. 5G), and reduction was exacerbated at P112 to a 79.5% decrease (P112: nested unpaired two-tailed t test, t= 3.630, df= 10, P= 0.0046, n= 23-26 hemisections and N= 6 hemicords from 3 mice per condition) (Fig. 5G-H). Spatial distributions of putative V1 Ia and Renshaw cells at P63 are shown in Fig. S4A-B. Altogether, these data show that all the analyzed V1 subpopulations are affected at symptomatic stage in the SOD1^G93A^ mice when compared to wt control. However, earlier in disease, V1 subpopulations show differential vulnerability, with Sp8+ neurons being more resistant to loss of transcripts, and putative Renshaw cells being most strongly affected.

**Fig. 5.**
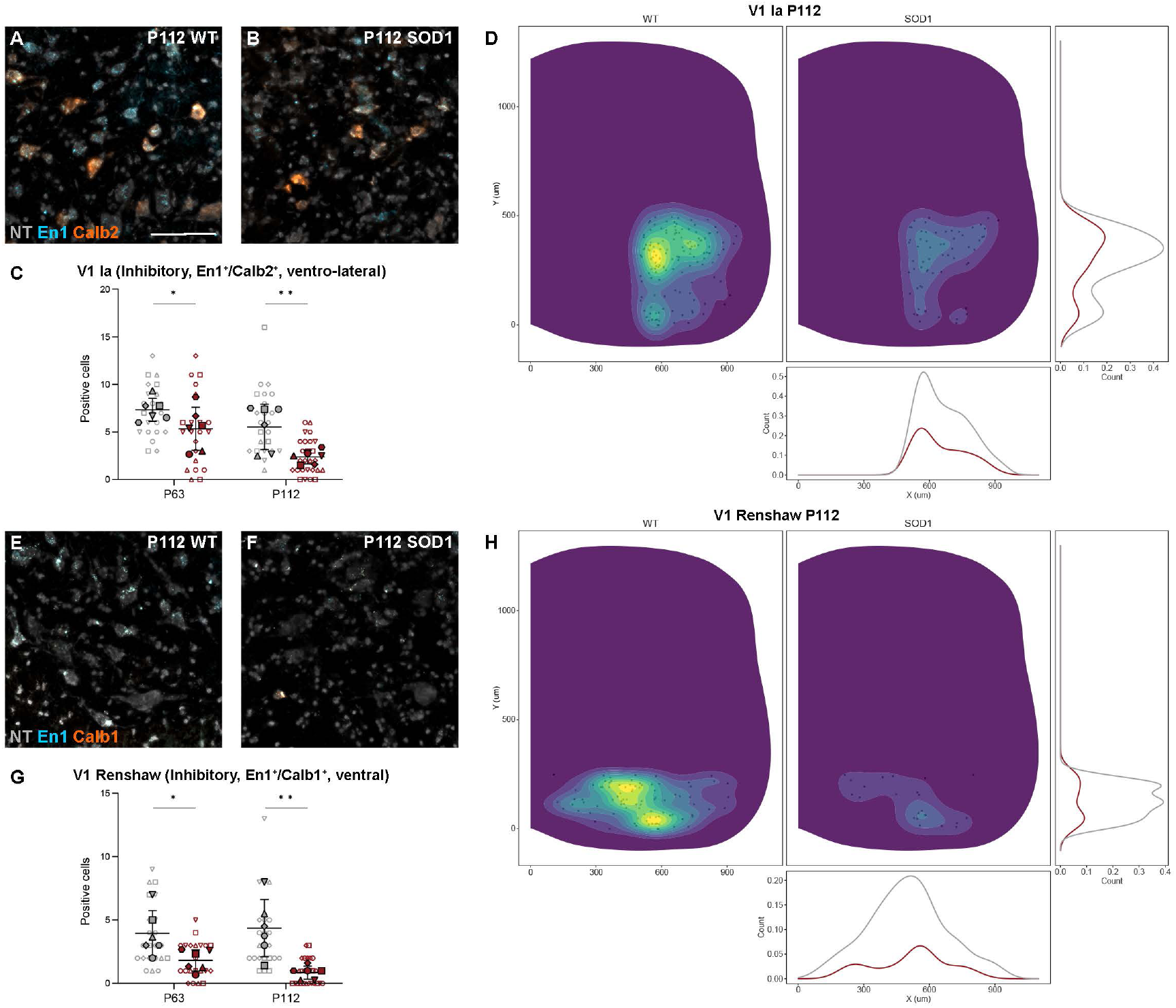
Putative V1 Ia interneurons and Renshaw cells show dysregulation in the SOD1^G93A^ mouse. (A-B) Co-localization of En1 (blue) and Calb2 (orange) transcript detected by RNAscope HiPlexUp in situ hybridization in wild-type (wt) (A) and SOD1^G93A^ mice (B) at P112. (C) Significant reduction in inhibitory En1+/Calb2+ neurons within the ventro-lateral region of the spinal cord, where V1 Ia interneurons are located, in SOD1^G93A^ mice (grey) versus wt (red) at P63 and P112 (nested unpaired two-tailed t tests; P63 P = 0.0498, P112 P = 0.0016). (D) Spatial distribution with count-based kernel density estimations of putative Ia interneurons from pooled P112 spinal cord sections. Data show the decrease in En1+/Calb2+ neurons in the SOD1^G93A^ mouse (right/red) compared to healthy controls (left/grey). (E-F) Co-localization of En1 (blue) and Calb1 (orange) transcript in wt (E) and SOD1^G93A^ spinal cord (F) at P112. (G) Dramatic loss of inhibitory En1+/Calb1+ neurons located within the ventral spinal cord, where Renshaw cells are found, in the SOD1^G93A^ mouse at P63, with exacerbation at P112 (nested unpaired two-tailed t tests; P63 P = 0.0271, P112 P = 0.0046). (H) Spatial distribution of putative Renshaw cells (En1+/Calb1+) at P112 with very few cells in the ventral spinal cord of SOD1^G93A^ mice compared to healthy control mice, evidencing the marked reduction in En1+/Calb1+ neurons. Scale bars, 100 μm. NT counterstaining in grey. 2D kernel densities in (D, H) were plotted with 10 bins using viridis scale. N = 6 hemicords from 3 mice (filled). Number of hemisections (empty): P30 n(wt) = 22, n(SOD1) = 20; P63 n(wt) = 22, n(SOD1) = 21; P112 n(wt) = 23, n(SOD1) = 26. Data shown as mean ± SD.

### Inhibitory neurotransmitter expression decreases only at symptomatic stages, at the same timepoint, survival of V1 interneurons is reduced

The expression of the glycine transporter 2 (Glyt2) and the enzymes catalyzing the GABA neurotransmitter Gad67 (also called Gad1) and Gad65 (also called Gad2) was investigated at the three timepoints (Fig. 6, S5, S6). These markers are used to visualize glycinergic and GABAergic neurons, respectively. GlyT2 transcript (Fig. 6A-B) was found downregulated only at P112 (P30: nested unpaired two-tailed t test, t= 0.332, df= 10, P= 0,7479, n= 22-20 hemisections and N= 6 hemicords from 3 mice per condition; P63: nested unpaired two-tailed t test, t= 0.881, df= 41, P= 0.3836, n= 22-21 hemisections and N= 6 hemicords from 3 mice per condition; P112: nested unpaired two-tailed t test, t= 3.058, df= 10, P= 0.0121, n= 23-26 hemisections and N= 6 hemicords from 3 mice per condition) (Fig. 6C-D). Gad67 expression (Fig. 6E-F) was also decreased only in symptomatic SOD1^G93A^ mice (P30: nested unpaired two-tailed t test, t= 0.105, df= 40, P= 0.9172, n= 22-20 hemisections and N= 6 hemicords from 3 mice per condition; P63: nested unpaired-two tailed t test, t= 1.604, df= 10, P= 0.1398, n= 22-21 hemisections and N= 6 hemicords from 3 mice per condition; P112: nested unpaired two-tailed t test, t= 3.399, df= 47, P= 0.0014, n= 23-26 hemisections and N= 6 hemicords from 3 mice per condition) (Fig. 6G-H), while no changes in Gad65 transcript levels (Fig. 6I-J) were observed at any of the investigated timepoints (P30: nested unpaired two-tailed t test, t= 0.414, df= 10, P= 0.6879, n= 22-20 hemisections and N= 6 hemicords from 3 mice per condition; P63: nested unpaired two-tailed t test, t= 1.273, df= 41, P= 0.2101, n= 22-21 hemisections and N= 6 hemicords from 3 mice per condition; P112: nested unpaired two-tailed t test, t= 0.995, df= 10, P= 0.3431, n= 23-26 hemisections and N= 6 hemicords from 3 mice per condition) (Fig. 6K-L).

**Fig. 6.**
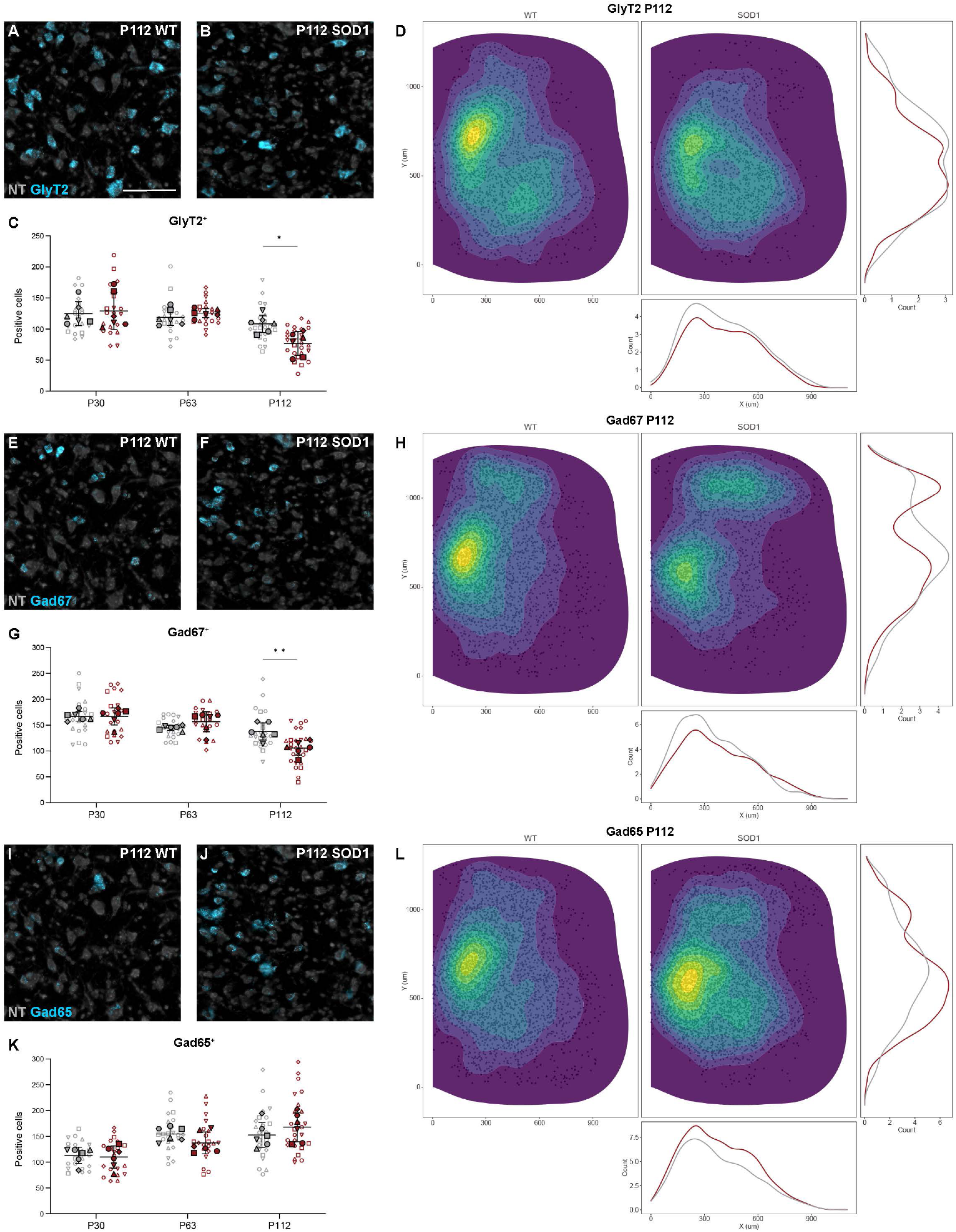
Inhibitory neurotransmitter markers show late affection in the SOD1^G93A^ mouse. (A-B, E-F, I-J) Microscopy images showing detection of transcript (blue) by RNAscope HiPlexUp in situ hybridization in healthy control mice and SOD1^G93A^ mice at P112 for GlyT2 (A-B), Gad67 (E-F), and Gad65 (I-J), with NT counterstaining (grey). (C, G, K) Quantification of positive cells for the different neurotransmitter markers in SOD1^G93A^ mice (red) versus healthy controls (grey). Data shows significant decrease in GlyT2+ neurons at P112 (C) (nested unpaired two-tailed t tests; P30 P = 0.7469, P63 P = 0.3836, P112 P = 0.0121), as well as Gad67+ cells at P112 (G) (nested unpaired two-tailed t tests; P30 P = 0.9172, P63 P = 0.1398, P112 P = 0.0014), but no changes in Gad65+ cells (K) (nested unpaired two-tailed t tests; P30 P = 0.6879, P63 P = 0.2101, P112 P = 0.3431). (D, H, L) Spatial distribution showing location and count-based kernel density estimations for GlyT2+ (D), Gad67+ (H) and Gad65+ (L) cells within the spinal cord of healthy control (left/red) and SOD1^G93A^ (right/grey) mice at P112. Scale bars, 100 μm. 2D kernel densities in (D, H, L) were plotted with 10 bins using viridis scale. N = 6 hemicords from 3 mice (filled). Number of hemisections (empty): P30 n(wt) = 22, n(SOD1) = 20; P63 n(wt) = 22, n(SOD1) = 21; P112 n(wt) = 23, n(SOD1) = 26. Data shown as mean ± SD.

To further investigate inhibitory interneuron survival, lineage tracing was performed for the V1 subpopulation in the SOD1^G93A^ mouse. Here, a triple transgenic mouse carrying the SOD1^G93A^ mutation and expressing tdTomato (39) under the En1 promoter (SOD1^G93A^;En1^Cre^;R26R^tdTomato^ mouse) was generated (Fig. 7A). TdTomato positive neurons were quantified in wt mice at different timepoints and compared with P112 SOD1^G93A^ mice (Fig. 7B-C). Quantifications revealed a decrease in number of tdTomato+ cells in the symptomatic mice (nested one-way ANOVA with Tukey’s multiple comparison; P63 wt tdTomato+ vs P112 SOD1 tdTomato+ P= 0.0006; wt n = 41 hemisections and N = 8 hemicords from 4 mice, SOD1 n = 27 hemisections and N = 6 hemicords from 3 mice) (Fig. 7D). We also investigated the presence of En1 transcript in tdTomato+ cells. This anlysis showed a parallel disappearance of the transcript (nested one-way ANOVA with Tukey’s multiple comparison; wt tdTomato+ vs tdTomato+/En1+ P = 0.8469, P112 SOD1 tdTomato+ vs tdTomato+/En1+ P = 0.7032; wt n = 41 hemisections and N = 8 hemicords from 4 mice, SOD1 n = 27 hemisections and N = 6 hemicords from 3 mice) (Fig. 7D). These results show that inhibitory interneurons die at later disease stage and that surviving V1 cells retain their transcript at late stages of the disease.

**Fig. 7.**
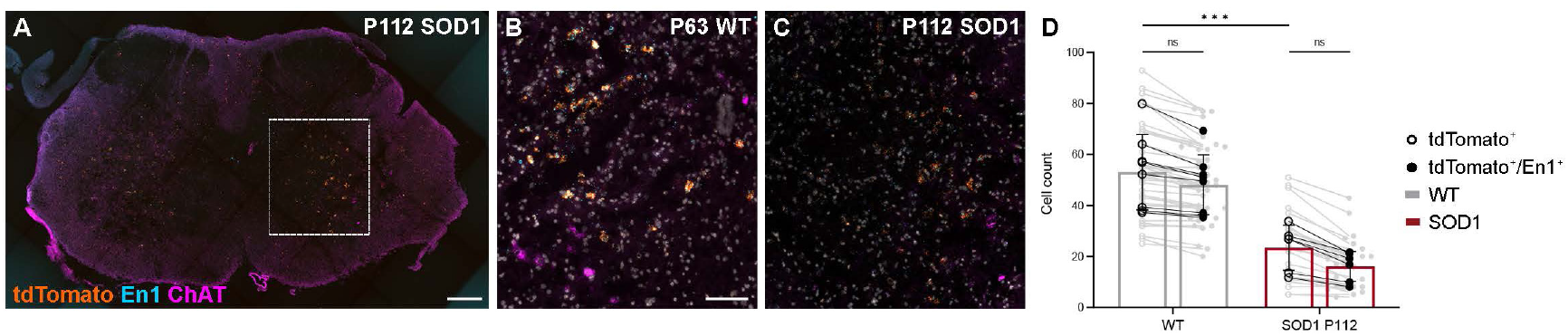
V1 interneuron fate in the SOD1^G93A^ mouse. (A) Microscopy image of a P112 SOD1^G93A^;En1^Cre^;R262R^tdTomato^ mouse lumbar spinal cord after RNAscope in situ hybridization, showing tdTomato (orange), En1 (blue), and ChAT (violet) transcript. (B-C) Magnification of tdTomato, En1 and ChAT transcript, with DAPI counterstaining (grey), in intermediate spinal cord of P63 En1^Cre^;R262R^tdTomato^ healthy control mice (wt) (B) and P112 SOD1^G93A^;En1^Cre^;R262R^tdTomato^ (SOD1) (C). (D) Quantification of tdTomato+ (empty) and tdTomato+/En1+ (filled) neurons in healthy control En1^Cre^;R262R^tdTomato^ (grey) and P112 SOD1^G93A^;En1^Cre^;R262R^tdTomato^ (red). Significant reduction of tdTomato+ cells was observed in the SOD1^G93A^ group versus control (nested one-way ANOVA with Tukey’s post hoc; wt tdTomato+ vs P112 SOD1 tdTomato+ P = 0.0006; wt n = 41 hemisections and N = 8 hemicords (from 4 mice), SOD1 n = 27 hemisections and N = 6 hemicords (from 3 mice)). No significant differences in tdTomato+ versus tdTomato+/En1+ were observed for either genotype (nested one-way ANOVA with Tukey’s post hoc; wt tdTomato+ vs tdTomato+/En1+ P = 0.8469, P112 SOD1 tdTomato+ vs tdTomato+/En1+ P = 0.7032; wt n = 41 hemisections and N = 8 hemicords (from 4 mice), SOD1 n = 27 hemisections and N = 6 hemicords (from 3 mice)), although SOD1^G93A^;En1^Cre^;R262R^tdTomato^ show a trend towards lower number of positive cells. Scale bars, 200 μm in (A) and 100 μm in (B-C). Data shown as mean ± SD.

### Glutamatergic V2a neurons lose Chx10 transcript later in disease, while Shox2+/Chx10- and V0G interneurons remain unaffected, and Vglut2 is not downregulated

Excitatory glutamatergic (Vglut2+) interneurons were also analyzed at the three different timepoints. Here, three main populations were investigated, the V2a (Chx10+), the Shox2+ (further divided in Chx10+ or Chx10-), and the V0_G_ (Pitx2+). V2a interneurons were defined as Chx10+ neurons found in the intermediate area of the spinal cord. When comparing transcript expression between wt and SOD1^G93A^ mice, a steep downregulation (57.6%) was observed only at P112 (P30: nested unpaired two-tailed t test, t= 1.392, df= 10, P= 0.1941, n= 22-20 hemisections and N= 6 hemicords from 3 mice per condition; P63: nested unpaired two-tailed t test, t= 0.110, df= 10, P= 0.9150, n= 22-21 hemisections and N= 6 hemicords from 3 mice per condition; P112: nested unpaired two-tailed t test, t= 4.894, df= 47, P < 0.0001, n= 23-26 hemisections and N= 6 hemicords from 3 mice per condition) (Fig. 8A-C). Analysis of spatial distribution at P112 revealed changes in both the ventral and dorsal areas (Fig. 8D), while no changes are observed at P30 and P63 (Fig. S7A-B).

**Fig. 8.**
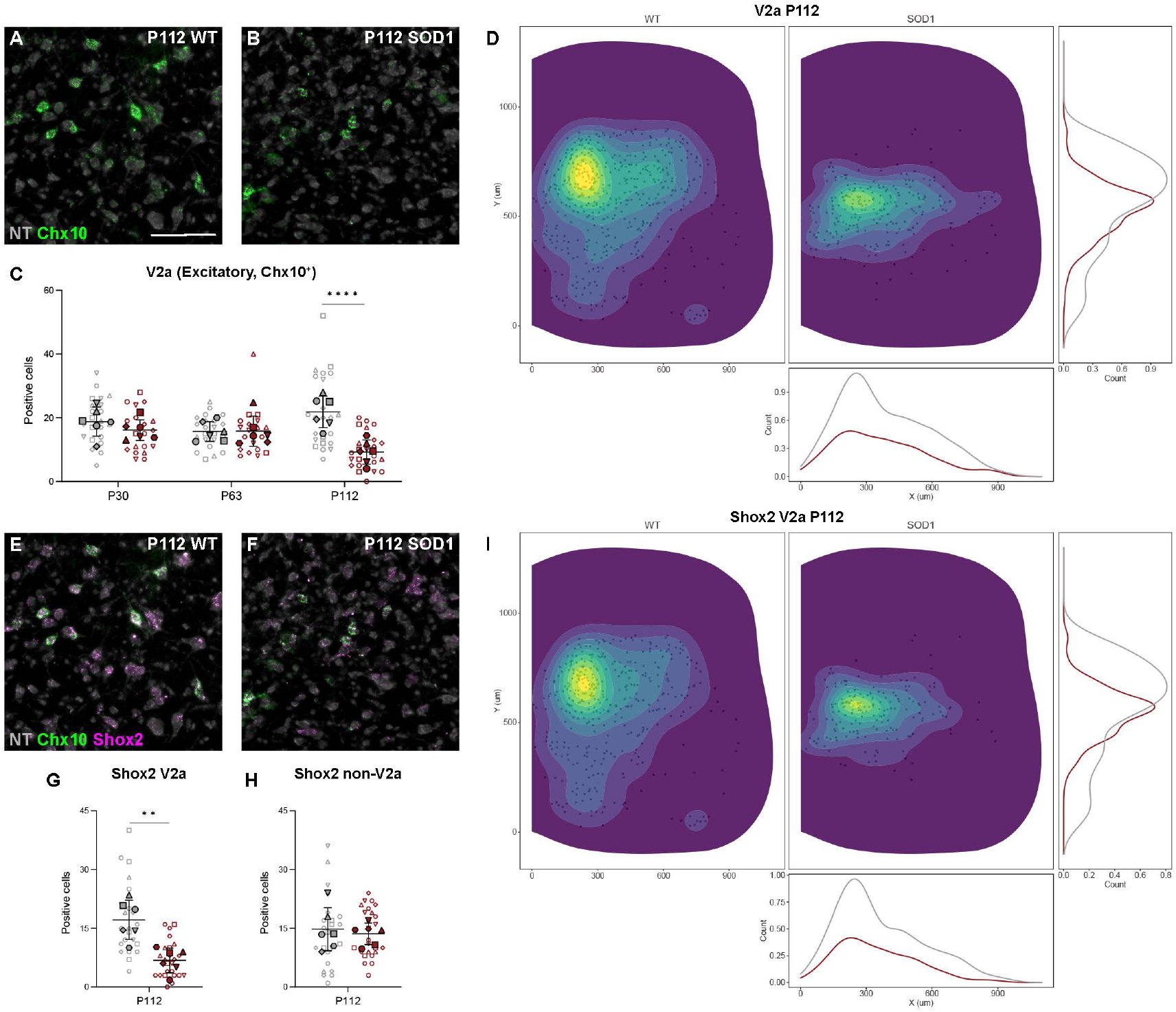
V2a and Shox2+-V2a interneurons are dysregulated at later stages of disease in the SOD1^G93A^ mouse. (A-B) RNAscope HiPlexUp in situ hybridization microscopy images showing detection of Chx10 transcript (green) in lumbar spinal cord of healthy control (wt) (A) and SOD1^G93A^ mice (B) at P112. (C) Significant reduction in number of excitatory (Vglut2+) Chx10+ neurons was revealed in SOD1^G93A^ mice (red) compared to healthy control mice (grey) only at P112 (nested unpaired two-tailed t tests; P30 P = 0.1941, P63 P = 0.9150, P112 P < 0.0001). (D) Spatial distribution of excitatory Chx10+ neurons with count-based kernel density estimations in SOD1^G93A^ mice (right/grey) versus healthy control mice (left/red) from pooled P112 sections show the loss of positive cells in the intermediate region of the spinal cord. (E-F) Co-localization of Chx10 (green) and Shox2 (violet) transcript by RNAscope HiPlexUp in wt (E) and SOD1^G93A^ spinal cord (F) at P112. (G-H) Quantification of excitatory Shox2+ neurons in combination with Chx10+ in SOD1^G93A^ mice versus healthy control mice at P112 revealed a steep decrease in Shox2+/Chx10+ neurons (G) but no changes in Shox2+/Chx10- (H) (nested unpaired two-tailed t tests; Shox2+ P = 0.0137, Shox2+/Chx10-P = 0.8914, Shox2+/Chx10+ P = 0013). (J) Spatial distribution of Shox2+/Chx10+ neurons within the spinal cord at P112 in SOD1^G93A^ mice versus healthy control mice shows a similar pattern to that observed for Chx10+ neurons. Scale bars, 100 μm. NT counterstaining in grey. 2D kernel densities in (D, J) were plotted with 10 bins using viridis scale. N = 6 hemicords from 3 mice (filled). Number of hemisections (empty): P30 n(wt) = 22, n(SOD1) = 20; P63 n(wt) = 22, n(SOD1) = 21; P112 n(wt) = 23, n(SOD1) = 26. Data shown as mean ± SD.

Shox2+ neurons can be divided in Shox2+/Chx10+ and Shox2+/Chx10- (16) (Fig. 8E-H). Here, we found that these two subpopulations are differentially affected in the SOD1^G93A^ mice. Interestingly, significant downregulation was observed among the double positive neurons at P112, with 59.6% reduction (P112: nested unpaired two-tailed t test, t= 4.410, df= 10, P= 0.0013, n= 23-26 hemisections and N= 6 hemicords from 3 mice per condition) (Fig. 8G). However, we did not observe changes in expression in the Shox2+/Chx10-population (P112: nested unpaired two-tailed t test, t= 0.137, df= 47, P= 0.8914, n= 23-26 hemisections and N= 6 hemicords from 3 mice per condition) (Fig. 8H). Changes in spatial distribution are shown in Fig. 8I and Fig. S7C-D.

Another glutamatergic population is the V0_G_ which comprises a small number of Pitx2 positive neurons located close to the central canal (7). The number of Pitx2 positive neurons was found unchanged at all timepoints, suggesting that this population remain unaffected during early and late stages in the SOD1^G93A^ mice (P30: nested unpaired two-tailed t test, t= 1.765, df= 40, P= 0.0852, n= 22-20 hemisections and N= 6 hemicords from 3 mice per condition; P63: nested unpaired two-tailed t test, t= 0.578, df= 10, P= 0.5762, n= 22-21 hemisections and N= 6 hemicords from 3 mice per condition; P112: nested unpaired two-tailed t test, t= 0.294, df= 10, P= 0.7746, n= 23-26 hemisections and N= 6 hemicords from 3 mice per condition) (Fig. S8A-D, Fig. S9B,D,F,H).

V2a (Chx10+), Shox2+ and V0_G_ (Pitx2+) neurons are all positive for the vesicular glutamate transporter 2 (Vglut2) (7, 16). We therefore also investigated Vglut2 expression at the three timepoints (Fig. 9). No changes were detected in Vglut2 expression at these stages (P30: nested unpaired two-tailed t test, t= 1.557, df= 40, P= 0.1273, n= 22-20 hemisections and N= 6 hemicords from 3 mice per condition; P63: nested unpaired two-tailed t test, t= 1.832, df= 30, P= 0.0742, n= 22-21 hemisections and N= 6 hemicords from 3 mice per condition; P112: nested unpaired two-tailed t test, t= 0.099, df= 10, P= 0.9225, n= 23-26 hemisections and N= 6 hemicords from 3 mice per condition) (Fig. 9C), nor was there a difference in the spatial distribution (Fig. 9D and Fig. S10). Altogether, these data suggest that some excitatory populations are affected later in disease and that, also in this case, specific markers are lost before the neurotransmitter. Moreover, the Chx10 and Shox2 populations show differential vulnerability, with Chx10+ and Shox2+/Chx10+ neurons being affected at P112, while the Shox2+/Chx10- and Pitx2+ neurons appear more resistant.

**Fig. 9.**
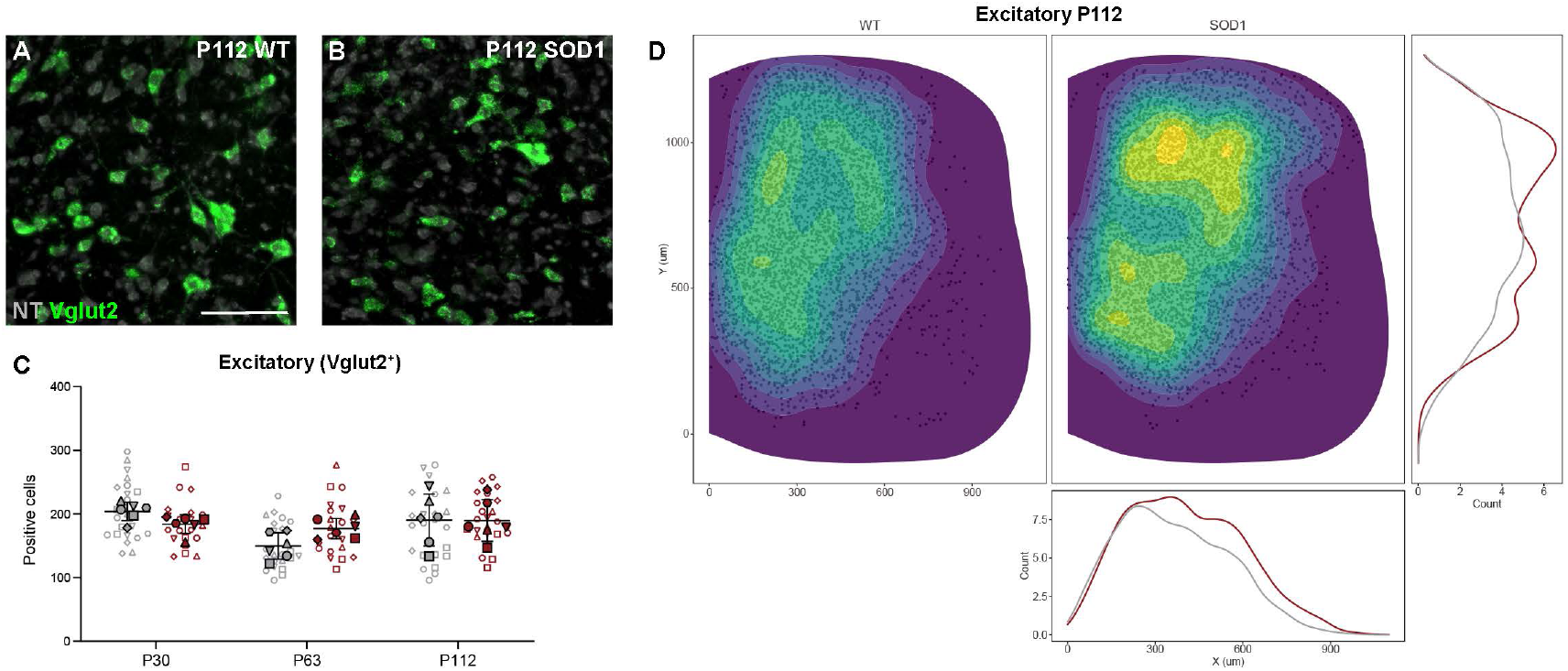
Expression of Vglut2 excitatory transcript shows no changes in the SOD1^G93A^ mouse. (A-B) Microscopy images after RNAscope HiPlexUp in situ hybridization showing Vglut2 transcript (green) within the lumbar spinal cord of wild-type (wt) (A) and SOD1^G93A^ mice (B) at P112, NT in grey. (C) Quantification of Vglut2+ neurons in SOD1^G93A^ mice versus healthy control mice revealed no changes at any of the tested timepoints (nested unpaired two-tailed t tests; P30 P = 0.1273, P63 P = 0.0742, P112 P = 0.9225). (D) Spatial distribution of Vglut2+ neurons within the spinal cord in healthy control mice (right/grey) and SOD1^G93A^ mice (left/red) at P112, including count-based kernel density estimations. The apparent higher density observed in the SOD1^G93A^ mouse is an effect of the higher number of sections used in this group. Scale bar, 100 μm. 2D kernel densities in (D) were plotted with 10 bins using viridis scale. N = 6 hemicords from 3 mice (filled). Number of hemi-sections (empty): P30 n(wt) = 22, n(SOD1) = 20; P63 n(wt) = 22, n(SOD1) = 21; P112 n(wt) = 23, n(SOD1) = 26. Data shown as mean ± SD.

### Cholinergic populations are affected at P112, except for V0C interneurons which do not lose the specific markers at any of the investigated timepoints

Spinal cholinergic neurons are heterogeneous (40), and in this study only somatic motor neurons and V0_C_ interneurons are investigated. The choline acetyltransferase (ChAT) marker is known to be downregulated upon muscle denervation (41), and motor neuron loss is significantly observed at P112 in the SOD1^G93A^ mouse model (26, 42, 43). We also found downregulation in ChAT+ cells located in the ventral spinal cord, putative motor neurons, at P112 (P112: nested unpaired two-tailed t test, t= 2.348, df= 47, P= 0.0231, n= 23-26 hemisections and N= 6 hemicords from 3 mice per condition), with 28.4% loss (Fig. S11). The V0_C_ interneurons, positive for Pitx2 and ChAT (7), were also analyzed (Fig. S8; Fig. S9 A,C,E,G). This population was not affected at these timepoints (P30: nested unpaired two-tailed t test, t= 0.257, df= 40, P= 0.7989, n= 22-20 hemisections and N= 6 hemicords from 3 mice per condition; P63: nested unpaired two-tailed t test, t= 0.578, df= 10, P= 0.6215, n= 22-21 hemisections and N= 6 hemicords from 3 mice per condition; P112: nested unpaired two-tailed t test, t= 0.123, df= 10, P= 0.6579, n= 23-26 hemisections and N= 6 hemicords from 3 mice per condition) (Fig. S9A,C,E,G). Thus, as expected, motor neurons are affected in the SOD1^G93A^ mice at symptomatic stage, while other cholinergic interneuron populations appear not to be.

## Discussion

Through this study, we have made significant progress in understanding the temporal dynamics of interneuron contribution to the progression of disease in SOD1^G93A^ mice. Our findings demonstrate that spinal interneurons exhibit differential susceptibility to the disease. Inhibitory interneurons exhibit earlier onset (starting at P63) but slower degeneration, while excitatory interneurons display later onset (starting at P112) but faster degeneration.

A condition for the success of this study was the development of methods to reliably detect co-expression of multiple markers in spinal cell populations. We developed two analysis pipelines that enable tracing of cell populations with high fidelity based on the detection and the quantification of multiple markers. With this approach we obtained a quantitative dataset on the relative changes of transcript expression in subpopulation of spinal interneurons throughout disease progression. Moreover, in this study we demonstrated that molecular markers that hitherto have been considered only expressed during development are indeed present into adult tissue and they can be used to delineate specific cell population derived from developmental progenitors. This finding allowed us to use these transcripts as cellular tracing markers also in the mouse adult spinal cord. Of particular interest are the V1 interneurons, given their early dysregulation and involvement in the SOD1^G93A^ locomotor phenotype (26). These novel results, together with our previous findings (26) suggest that in the SOD1^G93A^ mouse model V1 interneurons first lose connectivity with motor neurons, secondly are depleted of their specific markers and eventually of their neurotransmitter, before they die.

Furthermore, this study reveals distinct patterns of degeneration within different V1 interneuron populations. Putative Renshaw cells show the highest susceptibility to the disease, followed by putative Ia interneurons, the Foxp2 and Pou6f2 clades, and finally the Sp8 clade. While our analysis of Renshaw cell and Ia interneuron degeneration relies on putative definitions for these populations of interest, we assert that these neuronal populations are indeed involved, as the markers we employed are widely accepted within the field for identifying RCs and Ia neurons (11, 44, 45).

Notably, the observed changes in excitatory markers coincide with motor neuron death. This finding clearly demonstrate that inhibitory neurons dysregulation precede motor neuron death and thereby change the balance of the inhibitory excitatory synaptic input onto motor neurons. Therefore, pronounced motor neuron degeneration reflects a larger and consecutive impairment of spinal inhibitory and excitatory circuits in disease. Interestingly, inhibitory neurotransmitter deficits emerge at later stages of the disease compared to transcription factors. This implies the presence of a reservoir of transmitters earlier in disease, suggesting the maintenance of the inhibitory/excitatory phenotype of the neurons despite the ongoing degeneration and downregulation of transcrioption factor especially in the inhibitory populations. This aspect could be targeted as a potential ALS treatment aiming to maintain circuit connectivity (32) and restoring the excitatory/ inhibitory balance on motor neurons.

The observed degeneration of specific spinal interneuron subpopulations raises the key question of which mechanisms underlie their selective vulnerability. Although a definitive answer remains elusive, one possible framework, as suggested by Salamatina et al. (2020), proposes that interneurons with direct motor neuron connectivity are more prone to degeneration. This explanation aligns with much of the available data, such as the vulnerability of Renshaw cells and Ia interneurons compared to V1 clades or Shox2+ V2a interneurons compared to non-V2a interneurons. However, this framework contradicts the absence of degeneration in V0_C_ interneurons, which project onto motor neurons via C-boutons (7). Nevertheless, it is worth noting that V0_C_ interneurons constitute a small population of last-order neurons, which could account for their unique characteristics.

While this proposed framework is plausible, it does not directly clarify whether interneurons are the initial targets of degeneration, subsequently spreading the disease anterogradely to motor neurons, or if they degenerate due to a retrograde signal from motor neurons or because of connectivity loss. A strong indication of the role of inhibitory interneurons in disease initiation is the loss of V1 synaptic inputs onto motor neurons which precedes transcript downregulation and neuronal death. Thus, morphological evidence points towards an anterograde cascade starting from inhibitory interneurons, however gene discovery studies are needed to understand the molecular mechanisms behind this event. An alternative explanation for the reported vulnerability may lie in the differential connectivity to fast and slow motor neurons, which exhibit varying susceptibility. Once again, this notion aligns with the preferential vulnerability of V1 interneurons. However, it fails to explain the resistance observed in the V0_C_ subpopulation, as C-boutons have been shown to preferentially innervate fast motor neurons over slow ones (46, 47).

To gain further insights into the potential role of synaptic spread of the disease, it would be crucial to conduct a comprehensive analysis of the connectivity patterns between different interneuron populations and fast/slow motor neurons in both healthy and diseased conditions. Additionally, investigating the molecular mechanisms underlying the described synaptic loss could provide valuable information. These endeavors have the potential to shed light on the intricate mechanisms driving the selective vulnerability observed in spinal interneurons.

## Acknowledgments

We acknowledge the Department of Experimental Medicine at University of Copenhagen, especially Alex Soelberg Laugesen, and the Core Facility for Integrated Microscopy, Faculty of Health and Medical Sciences, University of Copenhagen. We thank Iryna Vesth-Hansen and Morten Bjerre for technical assistance, and Dr. Nicola Meola and Dr. Sara Wrobel at ACD Bio (Biotechne) for technical support and discussion.

Funding: This work was supported by the Lundbeck Foundation (I.A.), Louis-Hansen foundation (I.A.), the Læge Sofus Carl Emil Friis og hustru Olga Doris Friis foundation (I.A.), the Novo Nordisk Laureate Program ((O.K.), NNF15OC0014186), The Lundbeck Foundation ((O.K.) R345-2020-1769), the Louis-Hansen foundation (R.M.R.), The Faculty of Health and Medical Sciences at University of Copenhagen (O.K. and I.A.) and The School of Psychology and Neuroscience at University of St Andrews (I.A.).

## Author contributions

Conceptualization: IA, OK, RMR Methodology: IA, RMR, RS, PHV, JK Investigation: IA, RMR, RS, DBA Supervision: IA, OK Writing—original draft: IA, OK, RMR

## Data avalibility

The data that support the findings of this study is available from the corresponding authors upon reasonable request.

## Code avalibility

The code used to analyze data is available at https://github.com/Allodi-Lab/ISS_Pipeline and https://github.com/Allodi-Lab/HiPlexUp_Pipeline.

## Material and methods

### Experimental animals

Allanimal experiments were in a ccordance with the EUD irective 20110/63/EU and approved by the Danish Animal Inspectorate (Dyreforsøgstilsynet, ethical permit 2018-15-0201-01426) and the local ethics committee at the Faculty of Health and Medical Sciences. Wild-type C57BL6/J mice (Jackson Laboratory, strain #000664) were used for *in situ* sequencing experiments. Heterozygous SOD1^G93A^ mice (Jackson Laboratory, strain #004435 - B6.Cg-Tg(SOD1*G93A)1Gur/J) were used as ALS model. For V1 interneuron fate experiments, SOD1^G93A^ mice were crossed with heterozygous En1^Cre^ mice, a kind gift from Assistant Prof. Jay Bikoff (St. Jude Children’s hospital, St Louis, Texas, USA), and homozygous R26R^tdTomato^ (Jackson Laboratory, strain #007914 - B6.Cg-*Gt(ROSA)26Sortm14(CAG-tdTomato)Hze*/J). All strains and multiple transgenics were kept on a C57BL6/J genetic background. For experiments with SOD1^G93A^ mice, wild-type littermates were used as controls. Genotyping of SOD1^G93A^ was performed following supplier instructions. For quantification of human mutated SOD1 copy number by qPCR, Jackson’s SOD1^G93A^ founder breeder carrying 25 copies of the human mutated SOD1 gene was used as positive control, while a SOD1^127X^ carrying 19 copies was used as negative control. Mice were group-housed on a 12:12 h light:dark cycle with food and water *ad libitum*, 23-24 °C temperature, and 45-65% humidity.

### Preparation of fresh-frozen tissue

Samples were collected at postnatal days 1 and 28 for *in situ* sequencing experiments, postnatal days 30, 63 and 112 for RNAscope® HiPlexUp experiments, and postnatal days 63, 84 and 112 for V1 fate experiments. Mice were anesthetized with an overdose of pentobarbital solution (250 mg/kg) and sacrificed by decapitation. The lumbar region of the spinal cord was then quickly dissected and fresh frozen by immersion in dry ice cold isopentane for cryoprotection. For better preservation, the samples were stored at −80 °C and sectioned closely prior to use. Spinal cord coronal sections were embedded in Tissue-Tek® OCT Compound (Sakura, ref. 4583), cut on a cryostat at 12 μm (Thermo Fisher Scientific, Cryostar NX50 or Microm HM550), collected on SuperFrost Plus glass slides (Thermo Fisher Scientific), and further stored at −80 °C. To avoid RNA degradation, all equipment and tools involved in RNA work were cleaned with RNaseZap™ (Thermo Fisher Scientific, AM9780 or AM9786) prior to use.

### In situ sequencing

#### Assay

In situ sequencing was performed using CARTANA’s Library Prep Kit (CARTANA/10X Genomics, art. no. 1010-01/02; later replaced by Xenium, 10X Genomics). Sample preparation was performed as described in the user manual (doc. no. D023). All incubations were performed in SecureSealTM Hybridization Chambers (Grace Bio-Labs, ref. 621502). The used chimeric padlock probes were custom made by CARTANA and included the interneuron markers in Figure 1B. Quality control was performed using anchor probes labeled with Cy3 fluorophore to detect all rolling circle amplification products. Quality control images were taken in house with a 20x air objective using a Zeiss Axio Imager.Z1 or a Zeiss Axio Scan.Z1 slide scanner (EC Plan-Neofluar 20x/0.50 M27 or Plan-Apochromat 20x/0.8 M27, respectively; 0.227 μm/pixel) and approved by CARTANA/10X Genomics. Processed samples were then sent to CARTANA/10X Genomics for *in situ* barcode sequencing, imaging and data processing. Adapter probes and sequencing pools (containing AF488, Cy3, Cy5 and AF750 labels) were hybridized to the padlock probes to detect gene-specific barcodes through a sequence specific signal for each gene specific rolling circle amplification product. This was followed by imaging and performed 4 times in a row to allow for decoding of all barcoded probes. Raw data was obtained using a Nikon microscope system with 40x objective (0.165 μm/pixel) or CFI Plan-Apochromat λ 20x/0.75 (0.32 μm/pixel) and images included DAPI plus the four sequencing labels, taken as Z-stack and flattened using maximum intensity projection. After sequencing, the samples were sent back for further staining with NeuroTrace™ 530/615 or 640/660 (Invitrogen, N21482 or N21483) for segmentation purposes, which was performed by 2 h incubation at room temperature (1:200 in PBS).

#### Image and data processing

After image processing and decoding by CARTANA/10X Genomics, the data received included DAPI images of the sequenced spinal cord sections, and csv files with the gene information and coordinates in pixel units (0.32 μm/pixel) of the detected transcripts. Transcript data was presented in the format of low and high threshold, of which we used the low threshold. NeuroTrace™ imaging was performed using a Zeiss Axio Imager.Z1 or a Zeiss Axio Scan.Z1 slide scanner with EC Plan-Neofluar 20x/0.50 M27 or Plan-Apochromat 20x/0.8 M27, respectively (0.227 μm/pixel). DAPI counterstaining was also re-imaged together with the NeuroTrace™ for later image registration. Cell segmentation based on the NeuroTrace™ staining was performed using a custom deep learning model. The segmentation model was based on an ensemble of convolution neural networks (CNNs) trained on different partitions of the partial, manually annotated image data. Specifically, U-Net type architecture was used as the CNN configuration as the ensemble member. Predictions from the ensemble network yielded a probabilistic segmentation map for the presence of neurons. The cell boundary delineation was obtained using appropriate thresholds on these probabilistic segmentation masks. These cell segmentation results were registered to the sequencing data based on the NeuroTrace™ DAPI and the DAPI obtained during *in situ* sequencing assay. Keypoint-based registration using the random sample consensus (RANSAC) (48) algorithm was performed to obtain the transformation. The transformation obtained by registering the DAPIs was then applied to the NeuroTrace™ cell segmentation. These registered cells were then co-localized with the transcripts from CARTANA. Only transcripts that co-localized to segmented cells were considered. Further analysis was performed in RStudio (code at: https://github.com/Allodi-Lab/ISS_Pipeline).

### Quantitative multiplexed in situ hybridization

#### Assay

Quantitative multiplexed *in situ* hybridization was performed using the RNAscope® HiPlex12 Reagents (488, 550, 647) Kit Assay (Advanced Cell Diagnostics/Bio-Techne, cat. no. 324194; since replaced by HiPlex v2 Assay) in combination with the RNAscope® HiPlexUp Reagent (Advanced Cell Diagnostics/ Bio-Techne, cat. no. 324190). Sample pre-treatment and HiPlexUp assay were performed as described in the user manual (324100-UM) with the following modifications: fixation with 4% PFA was extended to 90 min during pre-treatment, samples were always stored in 5X Saline Sodium Citrate overnight at room temperature after probe hybridization, the concentration of Tween in the PBST was reduced to 0.05%, and the assay was adapted to the use of 3 fluorescent channels (4 rounds of fluorophores instead of 3). The combination of probes and fluorophores used, including catalogue numbers, is listed in Table 1. After the assay, samples were stained with NeuroTrace™ 530/615 (Invitrogen, N21482) by 2 h incubation at room temperature (1:200 in PBS) and re-imaged for segmentation purposes. Counterstaining with DAPI (provided in the kit) was performed as described in the manual and imaged in every round (including the NeuroTrace™ one) for image registration.

**Table 1.**
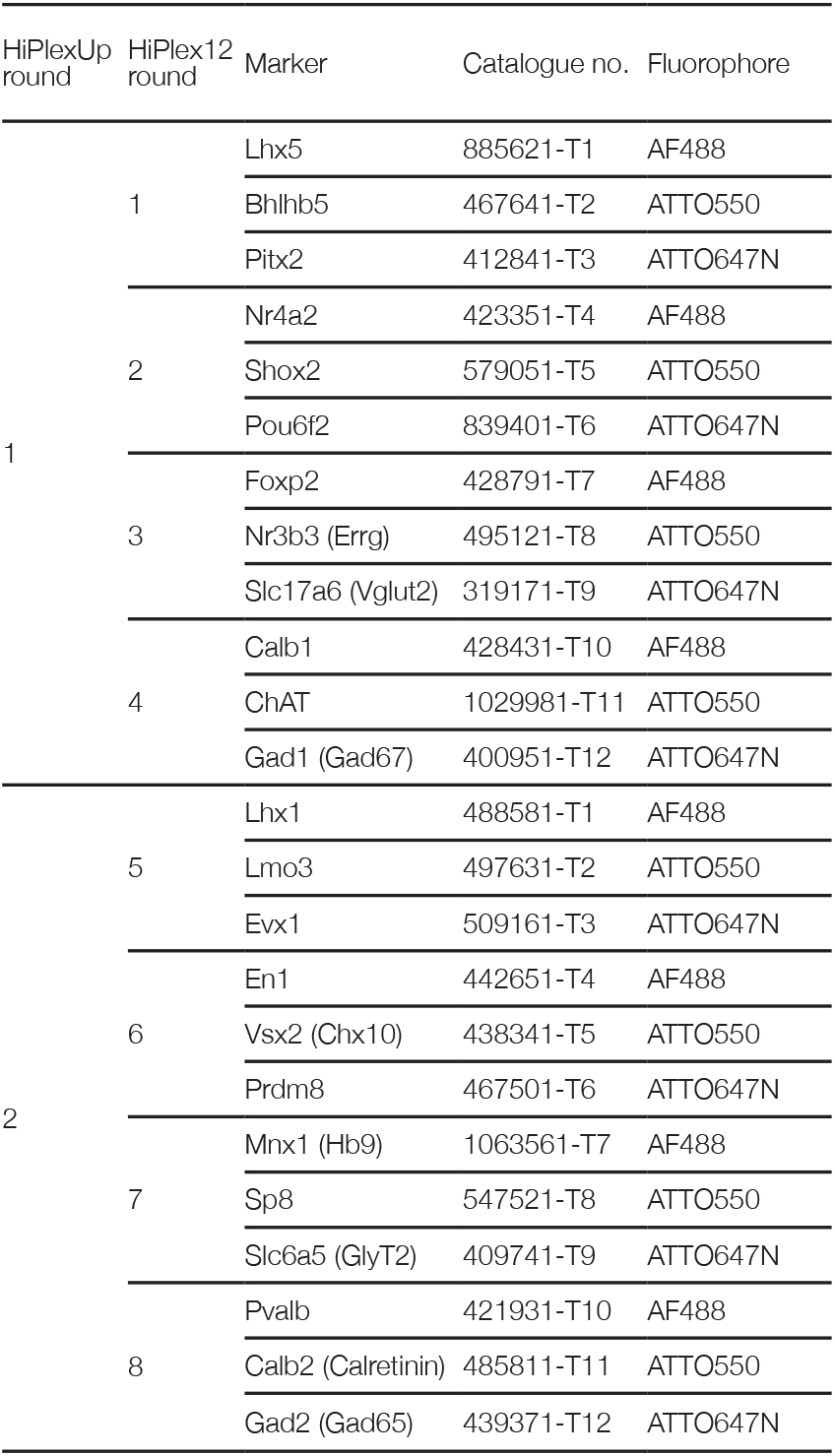
RNAscope® HiPlexUp probes. All probes were specific for mouse.

| HiPlexUp round | HiPlex12 round | Marker | Catalogue no. | Fluorophore |
| --- | --- | --- | --- | --- |
| 1 | 1 | Lhx5 | 885621-T1 | AF488 |
|  |  | Bhlhb5 | 467641-T2 | ATTO550 |
|  |  | Pitx2 | 412841-T3 | ATTO647N |
|  | 2 | Nr4a2 | 423351-T4 | AF488 |
|  |  | Shox2 | 579051-T5 | ATTO550 |
|  |  | Pou6f2 | 839401-T6 | ATTO647N |
|  | 3 | Foxp2 | 428791-T7 | AF488 |
|  |  | Nr3b3 (Errg) | 495121-T8 | ATTO550 |
|  |  | Slc17a6 (Vglut2) | 319171-T9 | ATTO647N |
|  | 4 | Calb1 | 428431-T10 | AF488 |
|  |  | ChAT | 1029981-T11 | ATTO550 |
|  |  | Gad1 (Gad67) | 400951-T12 | ATTO647N |
| 2 | 5 | Lhx1 | 488581-T1 | AF488 |
|  |  | Lmo3 | 497631-T2 | ATTO550 |
|  |  | Evx1 | 509161-T3 | ATTO647N |
|  | 6 | En1 | 442651-T4 | AF488 |
|  |  | Vsx2 (Chx10) | 438341-T5 | ATTO550 |
|  |  | Prdm8 | 467501-T6 | ATTO647N |
|  | 7 | Mnx1 (Hb9) | 1063561-T7 | AF488 |
|  |  | Sp8 | 547521-T8 | ATTO550 |
|  |  | Slc6a5 (GlyT2) | 409741-T9 | ATTO647N |
|  | 8 | Pvalb | 421931-T10 | AF488 |
|  |  | Calb2 (Calretinin) | 485811-T11 | ATTO550 |
|  |  | Gad2 (Gad65) | 439371-T12 | ATTO647N |

#### Image acquisition and processing

All imaging rounds were performed using a Zeiss Axio Scan.Z1 slide scanner. Samples from the same timepoints were prepared and imaged together. The same microscopy settings were used for all samples and imaging rounds. Full coronal spinal cord sections were scanned using a Plan-Apochromat 20x/0.8 M27 air objective (0.227 μm/pixel), with 3 Z-stack slices in increments of 2.5 μm and autofocused based on DAPI staining. A minimum of 5 sections, spaced 120 μm apart, were imaged per mouse – with 3 mice per condition and timepoint. Using the ZEN software (Zeiss, ZEN desk version 3.5) the resulting tiled images were first split by scene (each scene corresponding to a different section and imaging round – 728 HiPlexUp images + 91 NeuroTrace™ images), stitched based on DAPI staining, Z-stacks were flattened by maximum intensity projection, and single channel images were exported to TIF format keeping the original quality (3094 images). Sections that were too broken or folded were discarded at this point (at least 3 sections per mouse were kept). Images from different rounds corresponding to the same sections were then registered based on DAPI using the ‘Register Virtual Stack Slices’ and ‘Transform Virtual Stack Slices’ plugins in Fiji (ImageJ 2.3.0/1.53f51), and merged into single images (68 images) with 26 channels each (NeuroTrace™, DAPI, and the 24 interneuron markers). The merged images were converted back to CZI format for further analysis. To improve segmentation and reduce file sizes, images were cropped to include only the grey matter region of the spinal cord using the ZEN software ‘Extract polygons’ macro. These were then segmented based on NeuroTrace™ staining using the ZEN software Intellesis tool (80% confidence, split cells by watershed, 50 μm minimum size), and analyzed to obtain the coordinates and marker intensity profiles (including intensity mean, standard deviation and integrated intensity) for each detected cell. To detect positive cells, intensity thresholds were selected for each image and interneuron marker based on signal and background. Further analysis was performed in RStudio (see ‘Spatial profiling and quantifications’ section).

### Spatial profiling and quantifications

All coordinate normalizations, spatial profiling and positive cell quantifications for *in situ* sequencing and quantitative RNAscope®HiPlexUp *insitu* hybridizationwereperformedusing a custom analysis pipeline scripted in RStudio (2022.02.1 Build 461) (codes at: https://github.com/Allodi-Lab/HiPlexUp_Pipeline). Orientation angle, central canal coordinates, and maximum width and height of each hemisection were measured for coordinate normalization to a standardized mouse spinal cord hemisection. Spinal cord orientation was corrected using the ‘rearrr::rotate_2d’ function. Left and right hemisections were then split based on the coordinates of the central canal using the ‘dplyr::mutate’ function. Finally, coordinates were corrected for size and position using basic R functions. For *in situ* sequencing normalization, standardized full spinal cord: 2050 pixel (656 μm) medio-lateral, 2950 pixel (944 μm) dorsoventral in early post-natal (P1); 3450 pixel (1104 μm) mediolateral, 4400 pixel (1408 μm) dorso-ventral in young adult (P28). For *in situ* hybridization in adult, standardized grey matter only: 930 μm medio-lateral, 1280 μm dorso-ventral. The x-axis was defined as parallel to the medio-lateral axis with a medial origin, while the y-axis was defined as parallel to the dorso-ventral axis with a ventral origin. For *in situ* sequencing experiments, only transcripts that co-localized to segmented cells were visualized and considered for qualitative analysis. Co-localization of En1 transcript with other V1 markers was also visualized to represent V1 subpopulations. For RNAscope® HiPlexUp experiments, positive cells for each marker were identified based on the selected intensity thresholds using basic R functions, and combinations of markers with cell coordinates were used to describe and quantify putative interneuron populations. Spatial distributions show estimated counts and were calculated using a two-dimensional kernel density estimation with the ‘ggplot2::geom_density_2d_filled’ function and ‘count_var=“count”‘ argument.

### In situ hybridization

#### Assay

For assessment of V1 interneuron fate, *in situ* hybridization was performed using the RNAscope® Multiplex Fluorescent V2 Assay (Advanced Cell Diagnostics/Bio-Techne, cat. no. 323100). Sample pre-treatment and assay were performed as described for fresh frozen samples in the user manual (323100-USM), with overnight storage of samples in 5X Saline Sodium Citrate at room temperature after probe hybridization. The following probes (Advanced Cell Diagnostics/Bio-Techne) were used: Mm-En1-C1 (cat. no. 442651), tdTomato-C3 (cat. no. 317041-C3), and Mm-Chat-C2 as counterstaining (cat. no. 408731-C2). Opal™ 690, 570 and 520 dyes (Akoya Biosciences, FP1497001KT, FP1488001KT, FP147001KT) were used as fluorophores, respectively (1:1500 in TSA buffer, provided in the kit). DAPI was used as counterstaining (provided in the kit).

#### Image acquisition, processing and analysis

Images were acquired using a Zeiss Axio Scan.Z1 slide scanner with Plan-Apochromat 20x/0.8 M27 air objective (0.227 μm/pixel), with 5 Z-stack slices in increments of 1.5 μm and autofocused based on DAPI staining. A minimum of 5 sections (120 μm apart) were imaged per mouse – with 3 mice per condition and timepoint. Using the ZEN software (Zeiss, ZEN desk version 3.5) images were split by scene, stitched based on DAPI staining, Z-stacks were flattened by maximum intensity projection, and the ventral region of each hemisection was cropped (760 μm x 650 μm). These were then segmented based on tdTomato transcript signal using the ZEN software Intellesis tool (80% confidence, split cells by watershed, 50 μm minimum size), and analyzed to obtain the coordinates and intensity profiles. The number of segmented cells was used to quantify tdTomato+ cells, while further tdTomato+/En1+ quantification was based on En1 intensity thresholds.

### Statistical analysis

All statistical analyses were performed using the GraphPad Prism software (version 9.3.1(471)) unless otherwise stated. Sample sizes were not pre-determined by any statistical methods and were instead similar to previous publications (Allodi et al. 2021). For all experiments, mice were randomly allocated to different groups using a block design. Data collection was not blind to experimental group allocation but was semi-automated, analysis of data was performed blind. All mice used were included in the data sets. All RNAscope® HiPlexUp data were analyzed using a nested unpaired two-tailed t-test. Effect sizes were calculated as Cohen’s ds based on statistical data following the formula d_s_=t√((n_wt_+n_SOD1_)/(n_wt_n_SOD1_)) (where t is t-test value and n is the number of sections), and classified as trivial (< 0.2), small (0.2-0.49), medium (0.5-0.79) or large (≥0.8) as suggested in Cohen (1988). Reported means were calculated from the nested data. In figures, data are presented as individual values of replicates (empty shapes), mean value for each hemicord (filled shapes), and mean ± SD of hemicord values. The corresponding statistical values including n (and N), t, df, P values, mean difference ± SEM, 95% confidence interval of differences, and effect sizes are specified in the text, figure legends and tables S1-S3. V1 interneuron fate data was analyzed using nested one-way ANOVA with Tukey’s multiple comparisons test. Here data are presented as individual replicates (grey), mean value for each hemicord (black), and mean ± SD of hemicords. Statistical significance is reported as follows: *P < 0.05, **P < 0.01, ***P < 0.001, and ****P < 0.0001.

