## Supplementary information for "Spinal inhibitory neurons degenerate before motor neurons and excitatory neurons in a mouse model of ALS"

Montañana-Rosell et al. 2023

Contents:

- Figures S1 to S11
- Tables S1 to S3

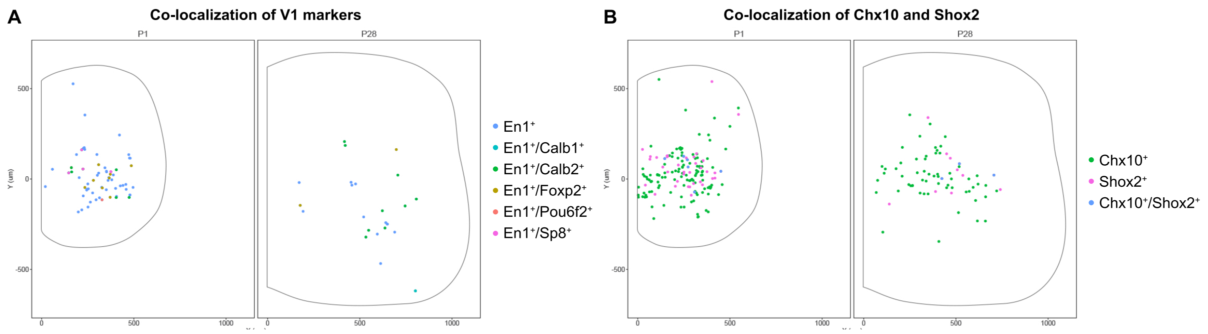


**Fig. S1. Co-localization of spinal interneuron identity markers with subpopulation markers.**

(A) Spatial distribution in early postnatal P1 (left) and young adult P28 (right) spinal cord of En1$^{+}$ cells (blue), as well as double positive cells where En1 co-localizes with V1 subpopulation and clade markers: En1$^{+}$/Calb1$^{+}$ (turquoise), En1$^{+}$/Calb2$^{+}$ (green), En1$^{+}$/Foxp2$^{+}$ (khaki), En1$^{+}$/Pou6f2$^{+}$ (salmon), and En1$^{+}$/Sp8$^{+}$ (pink).

(B) Spatial distribution of Chx10$^{+}$ (green) and Shox2$^{+}$ (pink), as well as double positive Chx10$^{+}$/Shox2$^{+}$ (blue) cells in the lumbar spinal cord at P1 (left) and P28 (right).

Data are pooled from n = 6 sections from N = 3 mice for P1; n = 5 sections from N = 5 mice for P28.

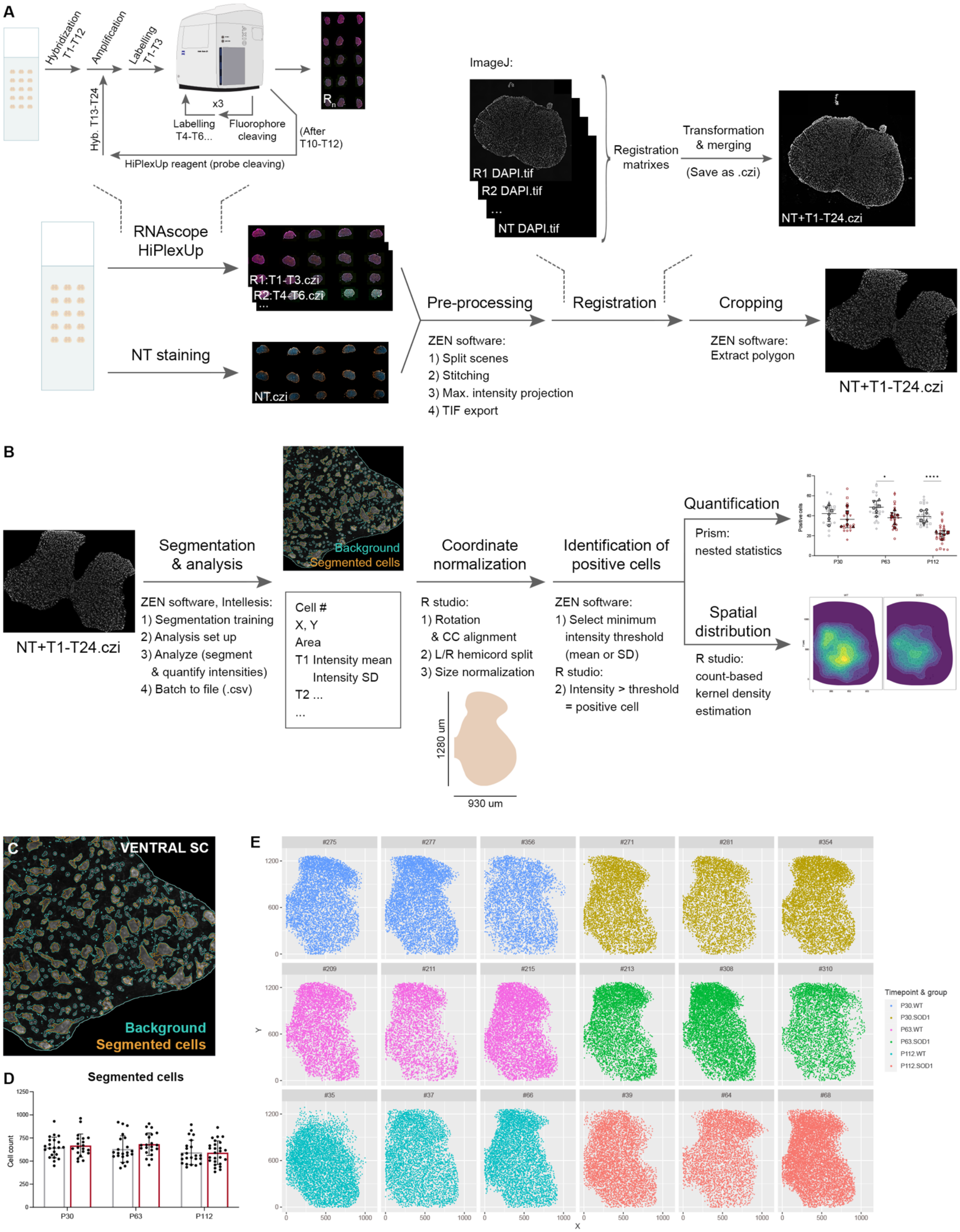


**Fig. S2. RNAscope HiPlexUp *in situ* hybridization analysis.**

(A) Workflow used for acquisition and pre-processing of multiplexing *in situ* hybridization data used in this study. *In situ* hybridization was performed using the RNAscope HiPlexUp assay, based on the sequential hybridization, amplification, labelling and imaging of probes. NeuroTrace (NT) staining together with DAPI was performed after RNAscope HiPlexUp assay for segmentation purposes during analysis. Obtained images were pre-processed with ZEN software as described. *In situ* hybridization images and NT, were then registered based on DAPI staining, transformed and merged using ImageJ. Lastly, spinal cord images were cropped in ZEN software to include the grey matter only. For each group and timepoint, a slide with spinal cords from 3 animals and 5-6 sections per animal were processed.

(B) Pipeline for analysis of RNAscope HiPlexUp data. Images obtained in (A) were segmented based on NT staining using the Intellesis tool from ZEN software. Segmented images were obtained together with quantitative data for each segmented cell (X, Y coordinates, areas and intensity profiles). Cell coordinates were then normalized to a standard spinal cord using a custom-made R script. Intensity thresholds were selected based on background and signal intensity and used to identify positive cells for each transcript. Combinatorial expression of transcripts together with coordinate restriction were used to define interneuron populations, which were quantified and plotted for analysis of interneuron dysregulation and spatial distribution.

(C) Example image of NT (grey) segmentation in the ventral region of the spinal cord, showing segmented cells (orange) and background (turquoise).

(D) Quantification of segmented cells in healthy control (wt) (grey) and $SOD1^{G93A}$ mice (red) at different timepoints showed no significant differences. Approximately 625 cells were segmented per hemisection on average. Data shown as mean ± SD, with individual hemisection values (black points).

(E) Spatial distribution of all segmented cells for each animal included in the study (all hemisections pooled). Each color represents a different group and timepoint: P30 wt in blue, P30 SOD1 in ocre, P63 wt in pink, P63 SOD1 in green, P112 wt in turquoise, P112 SOD1 in salmon.

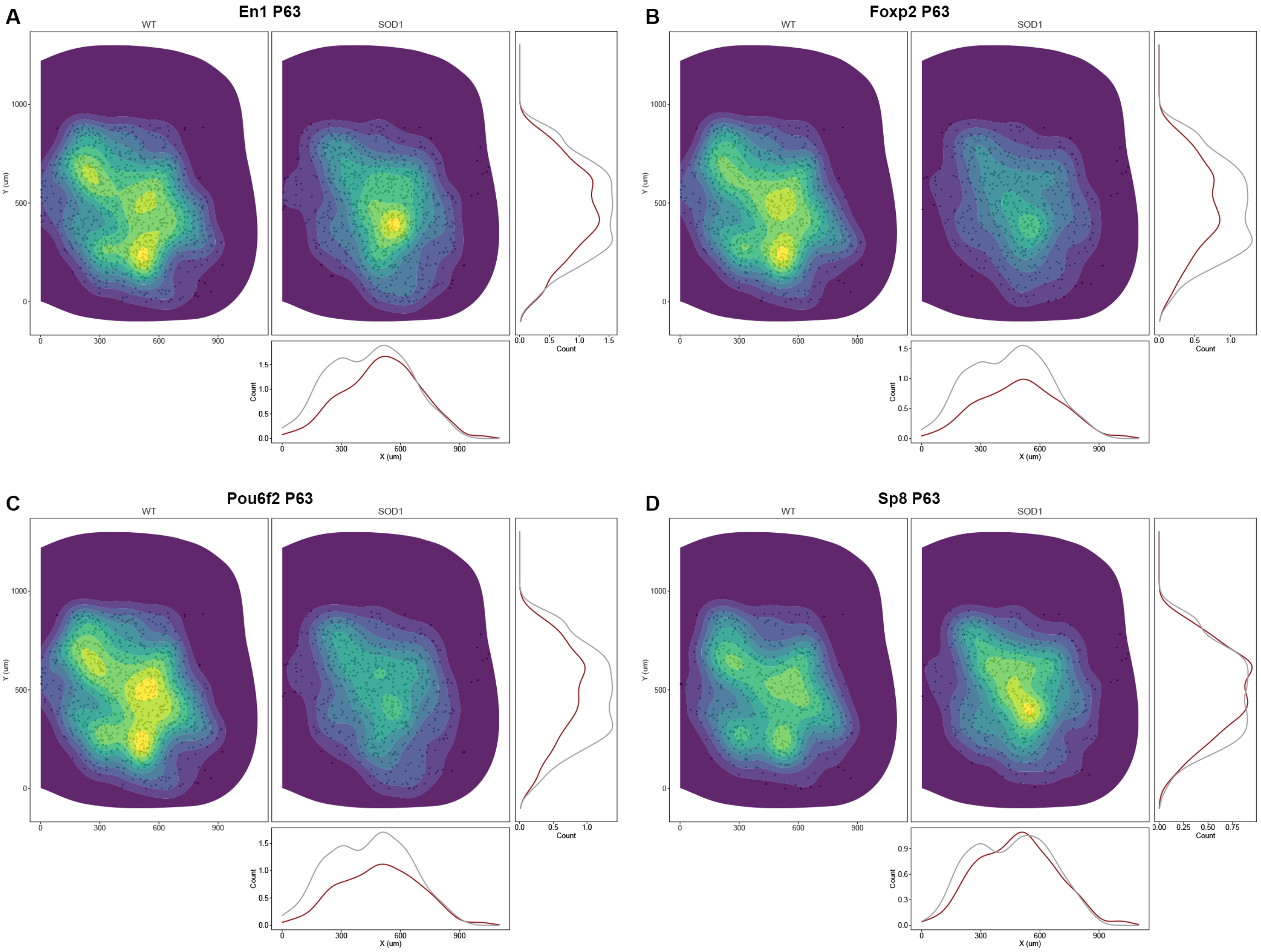


**Fig. S3. Spatial distribution of V1 interneurons and clades at P63.**

Spatial distribution with count-based kernel density estimations in 2D (main panel), X (bottom panel) and Y (right panel) dimensions in wild-type (wt) (left/grey) and SOD1$^{G93A}$ mice (right/red) at P63 for inhibitory En1$^+$ (A), En1$^+$/Foxp2$^+$ (B), En1$^+$/Pou6f2$^+$ (C) and En1$^+$/Sp8$^+$ (D) neurons. Loss of positive cells can be observed for overall En1$^+$ neurons, as well as Foxp2 and Pou6f2 clades.

Data were pooled from all P63 sections included in the study. 2D kernel densities were plotted with 10 bins using viridis scale. N = 6 hemicords from 3 mice, n(wt) = 22 hemisections, n(SOD1) = hemi21 sections.

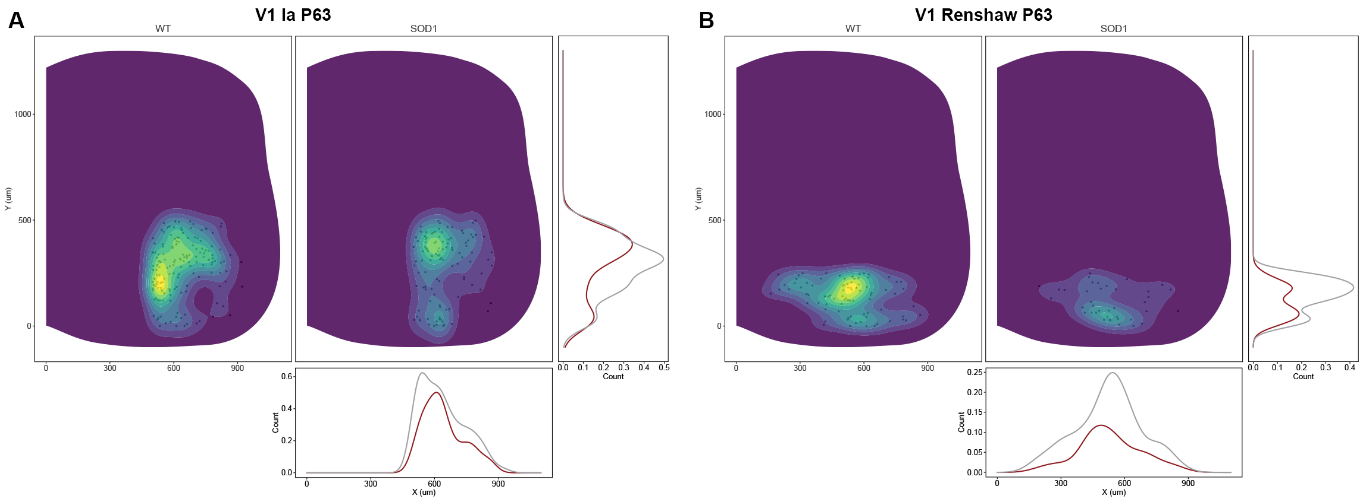


**Fig. S4. Spatial distribution of putative V1 Ia interneurons and putative Renshaw cells at P63.**

Spatial distribution with count-based kernel density estimations in 2D, X and Y dimensions of inhibitory $En1^+/Calb2^+$ neurons in the ventro-lateral region of the spinal cord (putative for V1 Ia interneurons) (A), and $En1^+/Calb1^+$ neurons in the ventral region of the spinal cord (putative for V1 Renshaw cells) (B) at P63. Comparison between healthy control (wt) (left/grey) and $SOD1^{G93A}$ mice (right/red) shows the reduction in positive cells for both putative populations in the $SOD1^{G93A}$ group at this timepoint, especially for putative Renshaw cells.

Data were pooled from all P63 sections included in the study. 2D kernel densities were plotted with 10 bins using viridis scale. N = 6 hemicords from 3 mice, n(wt) = 22 hemisections, n(SOD1) = 21 hemisections.

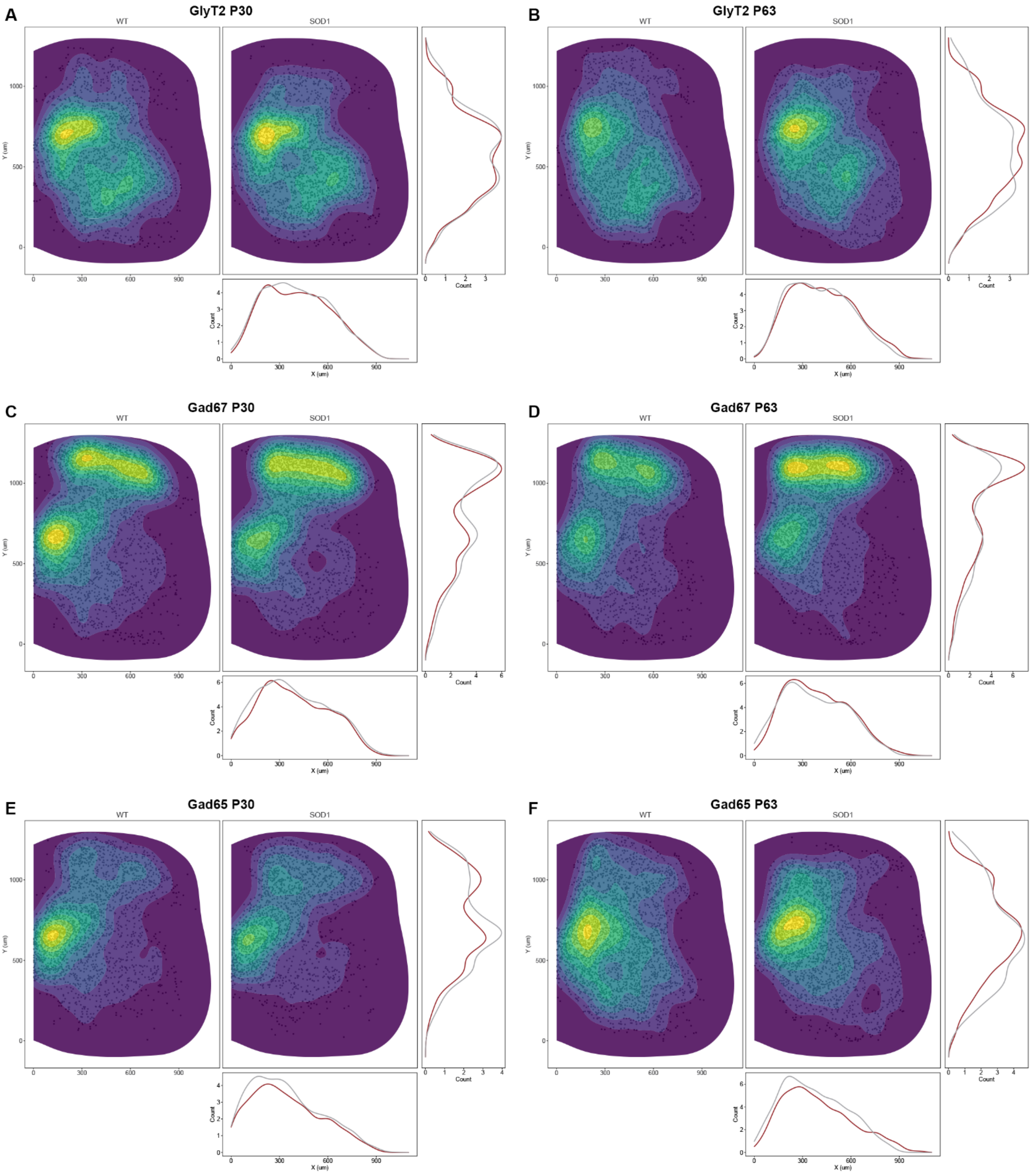


**Fig. S5. Spatial distribution of inhibitory neurotransmitter markers at P30 and P63.**

**(**A-B) Spatial distribution of GlyT2$^{+}$ neurons within the spinal cord of healthy control (wt) (left/grey) and SOD1$^{G93A}$ mice (right/red) at P30 (A) and P63 (B).

(C-D) Spatial distribution of Gad67$^{+}$ neurons at P30 (C) and P63 (D).

(E-F) Spatial distribution of Gad65$^{+}$ neurons at P30 (E) and P63 (F).

No differences in spatial distribution or positive cell densities were observed for any of the neurotransmitter markers at the two timepoints.

Data were pooled from all P30 or P63 sections included in the study. 2D kernel densities were plotted with 10 bins using viridis scale. N = 6 hemicords from 3 mice. Number of hemisections: P30 n(wt) = 22, n(SOD1) = 20; P63 n(wt) = 22, n(SOD1) = 21.

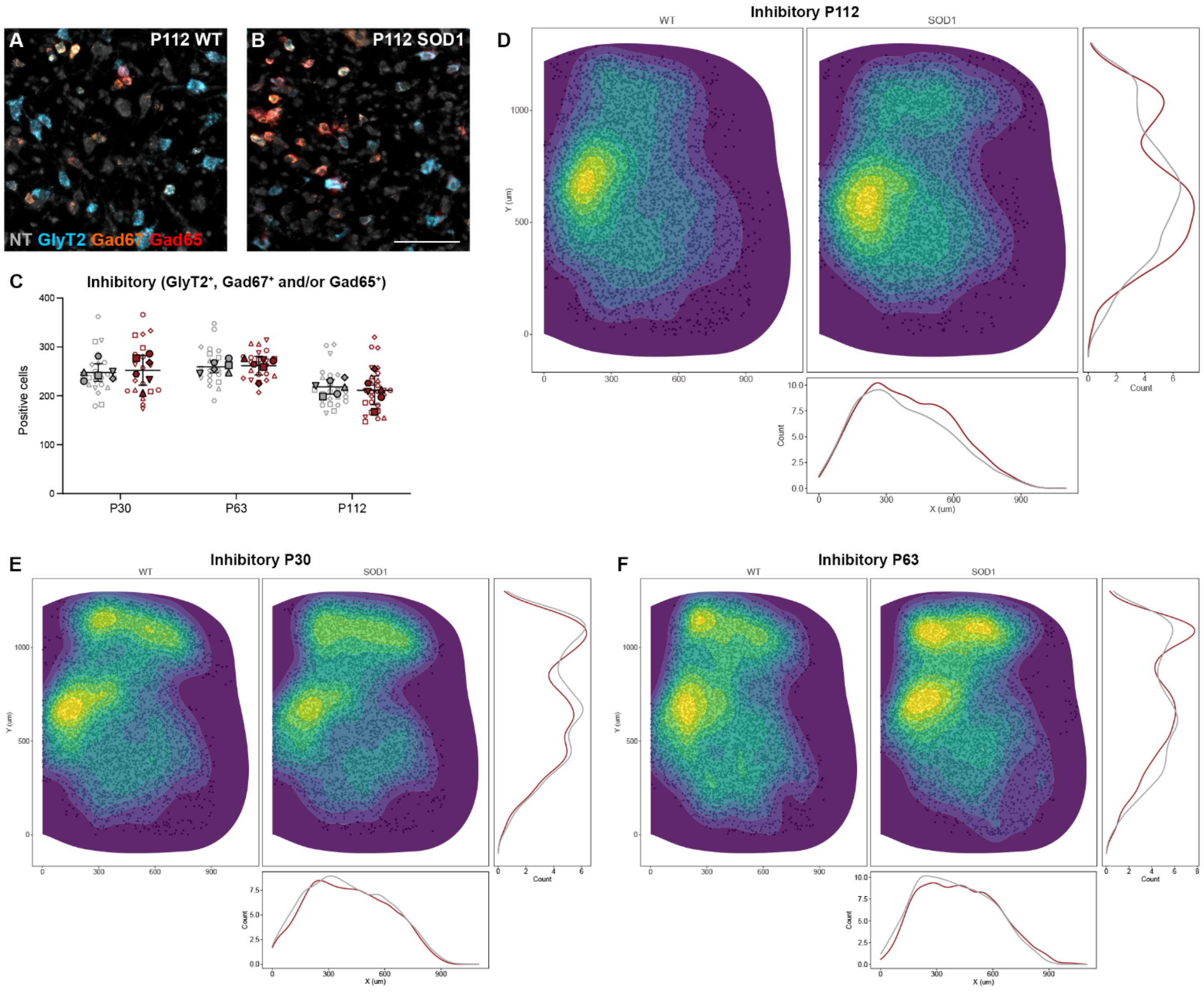


**Fig. S6. Joint expression of inhibitory neurotransmitter markers shows no changes in the SOD1$^{G93A}$ mouse.**

(A-B) Microscopy images of RNAscope HiPlexUp *in situ* hybridization showing GlyT2 (blue), Gad67 (orange) and Gad65 (red) transcript detection in healthy control (wt) (A) and SOD1$^{G93A}$ mice (B) at P112.

(C) Quantification of GlyT2$^{+}$, Gad67$^{+}$ and Gad65$^{+}$ neurons pooled together shows no differences at any of the assessed timepoints when comparing SOD1$^{G93A}$ (red) to healthy control mice (grey) (nested unpaired two-tailed t tests; P30 P = 0.6975, P63 P = 0.7634, P112 P = 0.7529).

(D-E) Spatial distribution including count-based kernel density estimations of pooled positive cells for the three inhibitory markers, in healthy control (left/grey) and SOD1$^{G93A}$ mice (right/red) at P112 (D) and P30 (E) and P63 (F).

Scale bar, 100 µm. 2D kernel densities in (D-F) were plotted with 10 bins using viridis scale. N = 6 hemicords from 3 mice (filled). Number of hemisections (empty): P30 n(wt) = 22, n(SOD1) = 20; P63 n(wt) = 22, n(SOD1) = 21; P112 n(wt) = 23, n(SOD1) = 26. Data shown as mean ± SD.

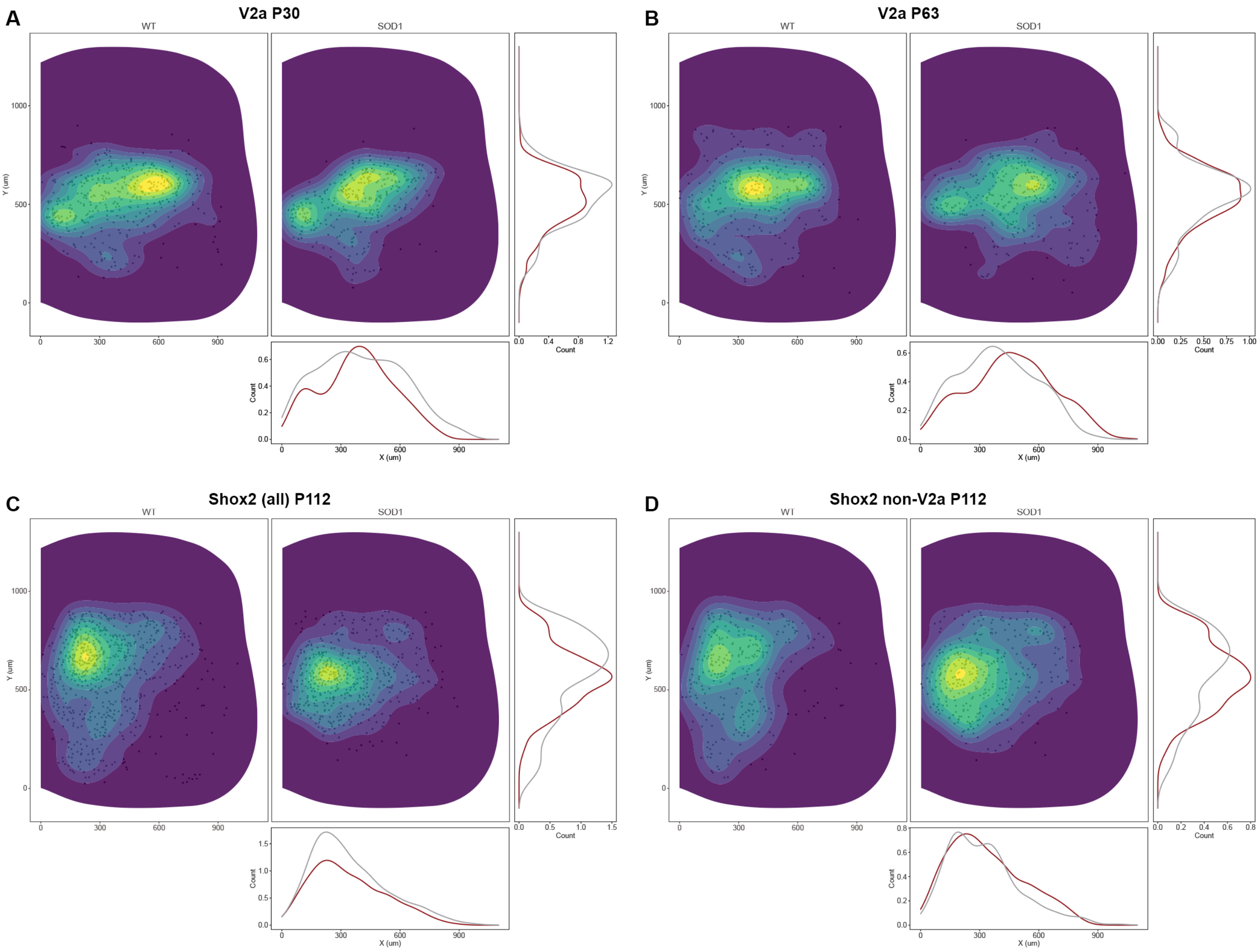


**Fig. S7. Spatial distribution of V2a interneurons at P30 and P63, and Shox2 and Shox2 non-v2a at P112.**

(A-B) Spatial distribution of excitatory Chx10$^{+}$ neurons within the spinal cord of healthy control (wt) (left/grey) and SOD1$^{G93A}$ mice (right/red) at P30 (A) and P63 (B), including count-based kernel density estimations. No obvious changes in density of positive neurons is seen between groups.

(C) Spatial distribution of all excitatory Shox2$^{+}$ neurons within the spinal cord at P112. Count-based kernel density estimations shows some loss of positive neurons in the SOD1$^{G93A}$ group compared to healthy control mice, in the intermediate region of the spinal cord.

(D) Spatial distribution of excitatory Shox2$^{+}$/Chx10$^{-}$ neurons at P112. The apparent slight increase of positive cells in SOD1$^{G93A}$ mice compared to healthy control mice (not observed upon quantification (Fig. 7)) is likely to be explained by the higher number of sections included in this group.

2D kernel densities were plotted with 10 bins using viridis scale. N = 6 hemicords from 3 mice. Number of hemisections: P30 n(wt) = 22, n(SOD1) = 20; P63 n(wt) = 22, n(SOD1) = 21; P112 n(wt) = 23, n(SOD1) = 26.

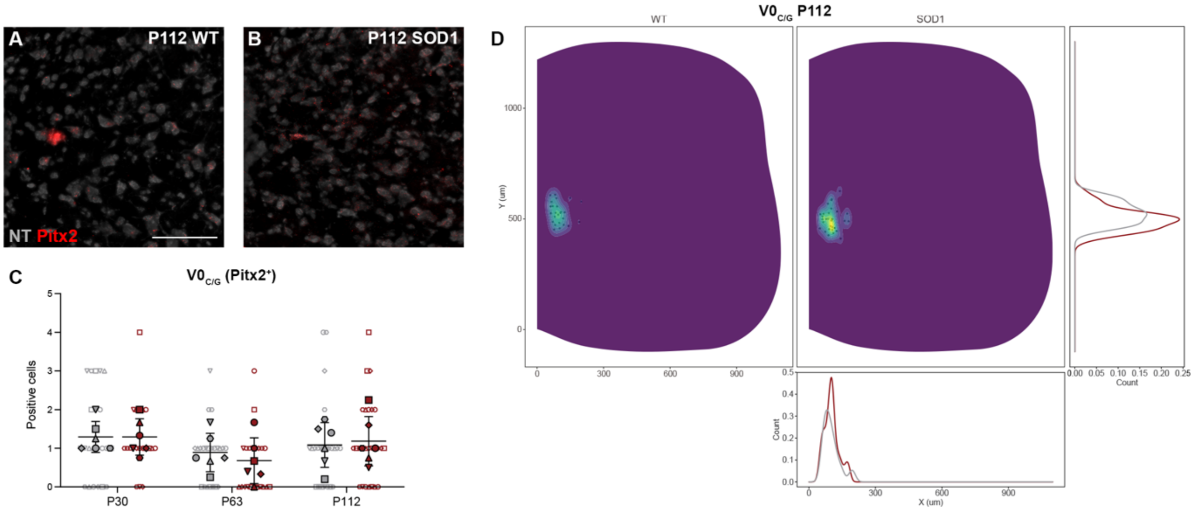


**Fig. S8. V0$_{C/G}$ interneuron transcript expression is not affected in the SOD1$^{G93A}$ mouse.**

(A-B) Imaging of Pitx2 transcript (red) detected by RNAscope HiPlexUp *in situ* hybridization in lumbar spinal cord of wild-type (wt) (A) and SOD1$^{G93A}$ mice (B) at P112, NT as counterstaining (grey).

(C) No changes were observed in number of Pitx2$^{+}$ neurons found around the central canal of SOD1$^{G93A}$ mice (red) compared to healthy control mice (red) at any of the tested timepoints (nested unpaired two-tailed t tests; P30 P = 0.8316, P63 P = 0.4776, P112 P = 0.7916).

(D) Spatial distribution including count-based kernel density estimations of Pitx2$^{+}$ neurons within the spinal cord of healthy control (right/grey) and SOD1$^{G93A}$ mice (left/red) at P112.

Scale bar, 100 µm. 2D kernel densities in (D) were plotted with 10 bins using viridis scale. N = 6 hemicords from 3 mice (filled). Number of hemisections (empty): P30 n(wt) = 22, n(SOD1) = 20; P63 n(wt) = 22, n(SOD1) = 21; P112 n(wt) = 23, n(SOD1) = 26. Data shown as mean ± SD.

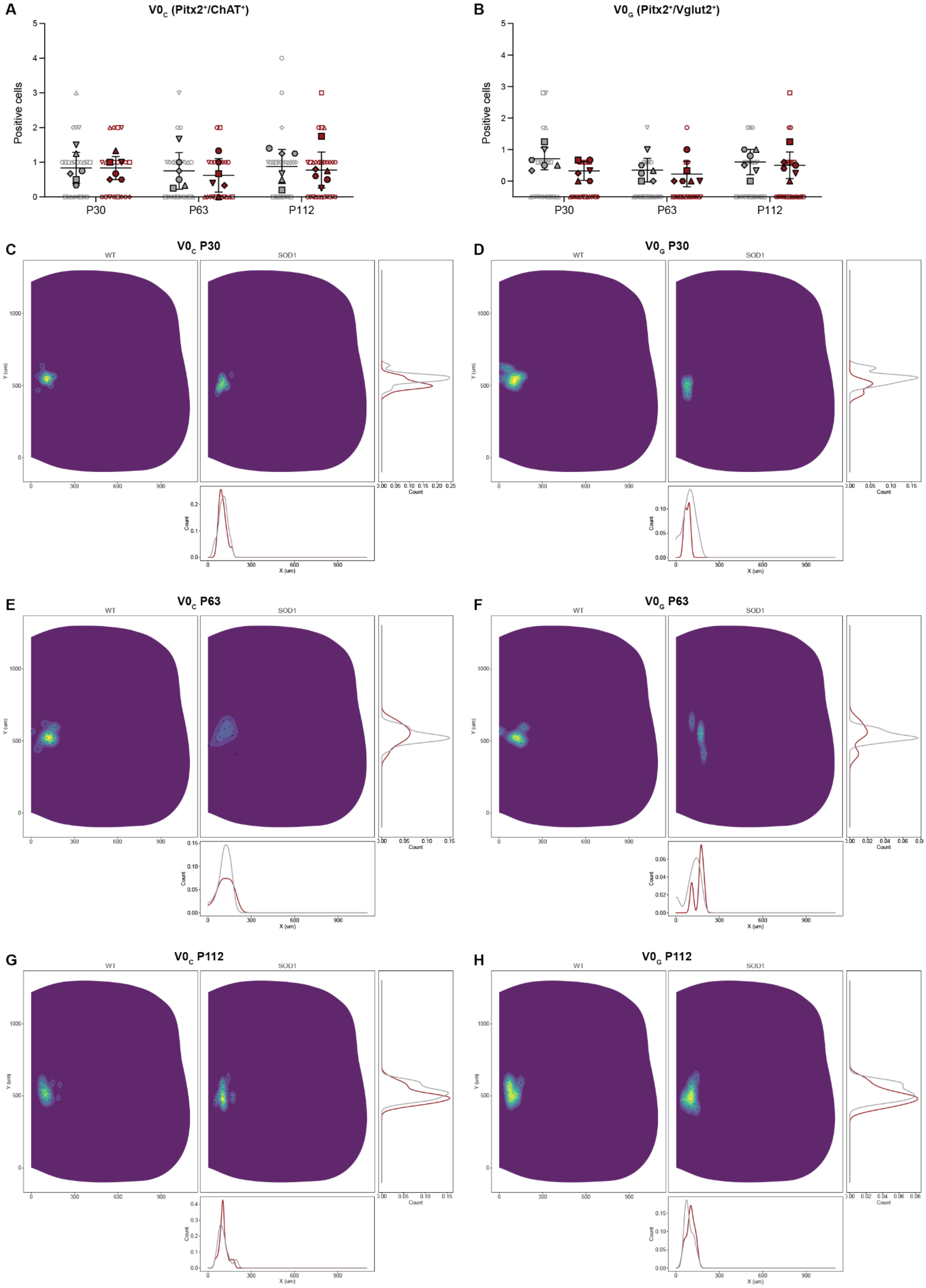
A
V0C (Pitx2+/ChAT+)
B
V0G (Pitx2+/Vglut2+)
Positive cells
P30
P63
P112
C
V0C P30
D
V0G P30
E
V0C P63
F
V0G P63
G
V0C P112
H
V0G P112
WT
SOD1
Y (um)
X (um)
Count

**Fig. S9. $V0_C$ and $V0_G$ show no transcript dysregulation in the $SOD1^{G93A}$ mouse.**

(A) Quantification of $Pitx2^+/ChAT^+$ neurons, putative for $V0_C$ interneurons, showed no differences between healthy control (grey) and $SOD1^{G93A}$ mice (red) at any of the analysed timepoints (nested unpaired two-tailed t tests; P30 P = 0.7989, P63 P = 0.6215, P112 P = 0.6579).

(B) Quantification of $Pitx2^+/Vglut2^+$ neurons, putative for $V0_G$ interneurons, also showed no differences between groups (nested unpaired two-tailed t tests; P30 P = 0.0852, P63 P = 0.5762, P112 P = 0.7746).

(C-H) Spatial distribution within the spinal cords of healthy control (left/grey) and $SOD1^{G93A}$ mice (right/red) at P30 (C-D), P63 (E-F), and P112 (G-H) for $Pitx2^+/ChAT^+$ neurons (C, E, G) and $Pitx2^+/Vglut2^+$ neurons (D, F, H), including count-based kernel density estimations.

2D kernel densities were plotted with 10 bins using viridis scale. N = 6 hemicords from 3 mice (filled). Number of hemisections (empty): P30 n(wt) = 22, n(SOD1) = 20; P63 n(wt) = 22, n(SOD1) = 21; P112 n(wt) = 23, n(SOD1) = 26. Data shown as mean ± SD.

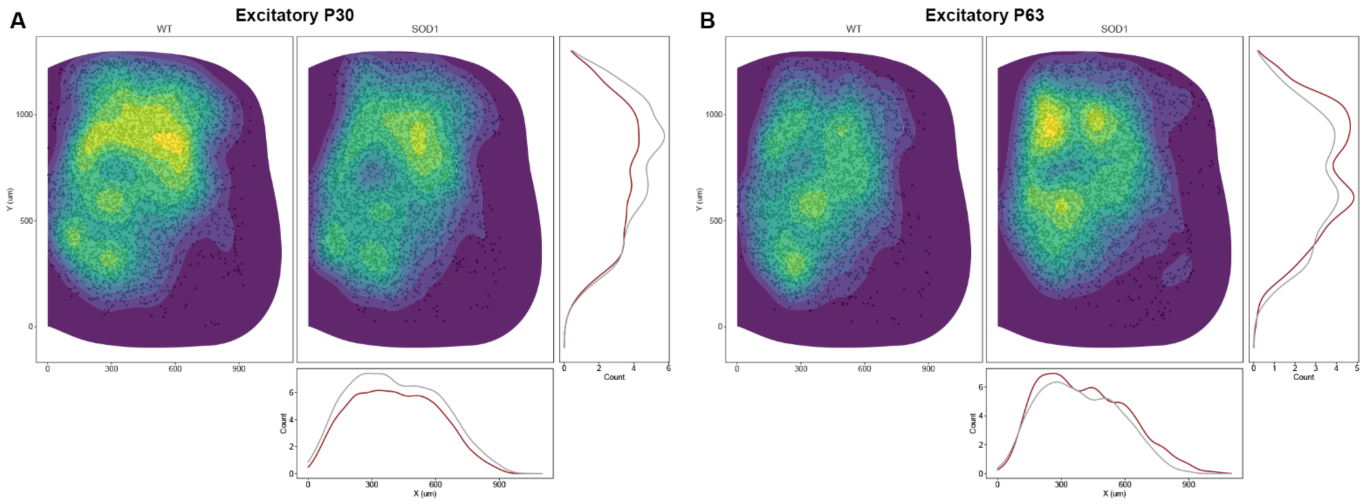


**Fig. S10. Spatial distribution of glutamatergic neurons at P30 and P63.**

Spatial distribution with count-based kernel density estimations of $Vglut2^+$ neurons within the lumbar spinal cord in healthy control (wt) (left/grey) and $SOD1^{G93A}$ mice (right/red) at P30 (A) and P63 (B).

2D kernel densities were plotted with 10 bins using viridis scale. N = 6 hemicords from 3 mice. Number of hemisections: P30 n(wt) = 22, n(SOD1) = 20; P63 n(wt) = 22, n(SOD1) = 21.

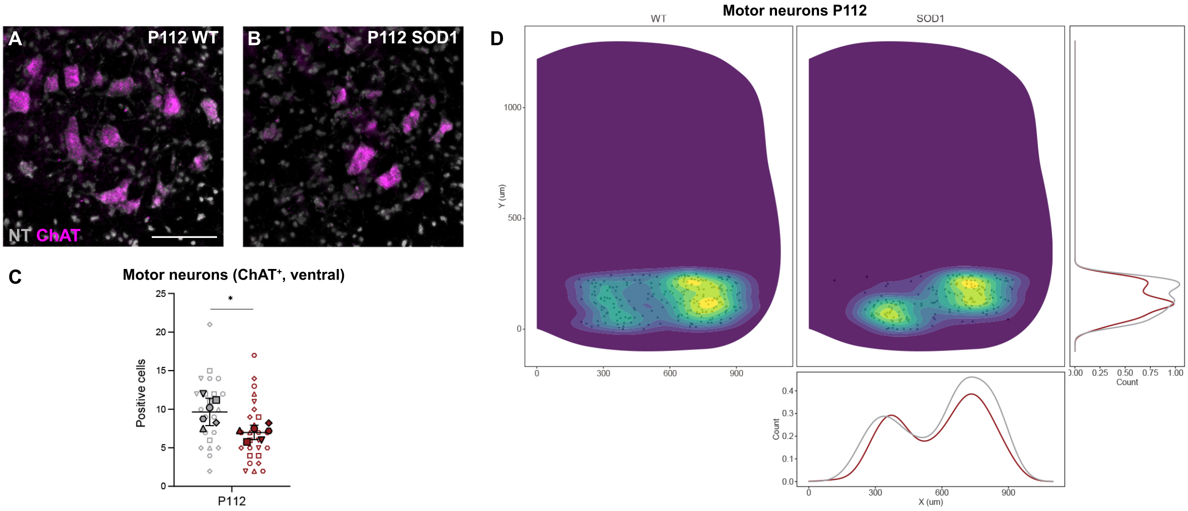


**Fig. S11. Quantification of ChAT transcript confirms motor neuron dysregulation at P112 in the SOD1$^{G93A}$ mouse.**

(A-B) Microscopy images showing ChAT transcript expression (violet) in motor neurons of the lumbar spinal cord of healthy control (wt) (A) and SOD1$^{G93A}$ mice (B) at P112, NT counterstaining (grey).

(C) Quantification of ChAT$^{+}$ neurons in the ventral region of the spinal cord, where motor neurons are located, showed significant reduction in the SOD1$^{G93A}$ group (red) compared to healthy control mice (grey) at P112 (nested unpaired two-tailed t tests; P112 P = 0.0231).

(D) Spatial distribution of ChAT$^{+}$ neurons within the ventral spinal cord in healthy control (right/grey) and SOD1$^{G93A}$ mice (left/red) at P112, shown with count-based kernel density estimations.

Scale bar, 100 µm. 2D kernel densities in (D) were plotted with 10 bins using viridis scale. N = 6 hemicords from 3 mice (filled). Number of hemisections (empty): P112 n(wt) = 23, n(SOD1) = 26. Data shown as mean ± SD.

**Table S1. Sample and replicate numbers.** Number of replicates included for each timepoint and condition included in the study, including biological (hemicords/mice, N) and technical replicates (hemisections, n). The number of sections per hemicord and the shapes used to represent them in the figures are also specified.

| Timepoint | Group | Hemicords (N) | Sections (n) | Sections/hemicord |
|---|---|---|---|---|
| P30 | wt | 6 (3 mice) | 22 | ● 4 ■ 4<br>▲ 4 ▼ 4<br>◆ 3 ⬣ 3 |
| | SOD1 | 6 (3 mice) | 20 | ● 3 ■ 3<br>▲ 3 ▼ 3<br>◆ 4 ⬣ 4 |
| P63 | wt | 6 (3 mice) | 22 | ● 4 ■ 4<br>▲ 3 ▼ 3<br>◆ 4 ⬣ 4 |
| | SOD1 | 6 (3 mice) | 21 | ● 3 ■ 3<br>▲ 4 ▼ 5<br>◆ 3 ⬣ 3 |
| P112 | wt | 6 (3 mice) | 23 | ● 5 ■ 5<br>▲ 2 ▼ 3<br>◆ 4 ⬣ 4 |
| | SOD1 | 6 (3 mice) | 26 | ● 4 ■ 4<br>▲ 4 ▼ 4<br>◆ 5 ⬣ 5 |

**Table S2. Statistical details for assessment of inhibitory populations.**

| Marker/s (Population) [Figure] | Time-point | n sections (N = 3 mice, 6 hemicords) | | Nested unpaired two-tailed t-test | | | wt mean | SOD1 mean | Descriptive report of differences | | Effect size (d) | Est. % loss |
|---|---|---|---|---|---|---|---|---|---|---|---|---|
| | | wt | SOD1 | t | df | P value | | | (SOD1-wt) ±SEM | 95% CI of diff. | | |
| Inhibitory/En1+ (V1) [Fig. 3A-D] | P30 | 22 | 20 | 1.305 | 10 | 0.2210 (ns) | 42.31 | 36.13 | -6.179 ± 4.733 | -16.73 to 4.367 | 0.403 (small) | - |
| | P63 | 22 | 21 | 2.493 | 10 | 0.0318 (*) | 47.92 | 38.74 | -9.182 ± 3.682 | -17.39 to -0.977 | 0.761 (med.) | 19.16 |
| | P112 | 23 | 26 | 6.379 | 47 | <0.0001 (****) | 40.04 | 21.88 | -18.16 ± 2.847 | -23.89 to -12.43 | 1.826 (large) | 45.35 |
| Inh./En1+/Foxp2+ (V1 Foxp2 clade) [Fig. 3E-F, K] | P63 | 22 | 21 | 3.777 | 10 | 0.0036 (**) | 38.58 | 24.14 | -14.44 ± 3.824 | -22.96 to -5.921 | 1.152 (large) | 37.43 |
| | P112 | 23 | 26 | 5.502 | 47 | <0.0001 (****) | 33.78 | 18.85 | -14.95 ± 2.715 | -20.40 to -9.475 | 1.575 (large) | 44.26 |
| Inh./En1+/Pou6f2+ (V1 Pou6f2 clade) [Fig. 3G-H, L] | P63 | 22 | 21 | 3.168 | 10 | 0.0100 (*) | 42.88 | 27.84 | -15.03 ± 4.746 | -25.61 to -4.459 | 0.966 (large) | 35.05 |
| | P112 | 23 | 26 | 3.637 | 47 | 0.0007 (***) | 25.70 | 14.42 | -11.27 ± 3.100 | -17.51 to -5.037 | 1.041 (large) | 43.85 |
| Inh./En1+/Sp8+ (V1 Sp8 clade) [Fig. 3I-J, M] | P63 | 22 | 21 | 0.328 | 41 | 0.7444 (ns) | 27.36 | 26.10 | -1.268 ± 3.865 | -9.073 to 6.536 | 0.100 (trivial) | - |
| | P112 | 23 | 26 | 2.754 | 47 | 0.0083 (**) | 22.43 | 14.27 | -8.166 ± 2.965 | -14.13 to -2.201 | 0.788 (med.) | 36.41 |
| Inh./En1+/Calb2+ (putative V1 Ia) [Fig. 4A-D] | P63 | 22 | 21 | 2.022 | 41 | 0.0498 (*) | 7.273 | 5.238 | -2.035 ± 1.006 | -4.067 to -0.00216 | 0.617 (med.) | 27.98 |
| | P112 | 23 | 26 | 4.291 | 10 | 0.0016 (**) | 6.012 | 2.383 | -3.629 ± 0.8459 | -5.514 to -1.745 | 1.228 (large) | 60.36 |
| Inh./En1+/Calb1+ (putative V1 Renshaw) [Fig. 4E-H] | P63 | 22 | 21 | 2.587 | 10 | 0.0271 (*) | 3.876 | 1.830 | -2.046 ± 0.7908 | -3.808 to -0.284 | 0.789 (med.) | 52.79 |
| | P112 | 23 | 26 | 3.630 | 10 | 0.0046 (**) | 4.203 | 0.860 | -3.343 ± 0.9208 | -5.395 to -1.291 | 1.039 (large) | 79.54 |
| GlyT2+ (Glycinergic) [Fig. 6A-D] | P30 | 22 | 20 | 0.332 | 10 | 0.7469 (ns) | 123.9 | 128.5 | 4.620 ± 13.92 | -26.40 to 35.64 | 0.103 (trivial) | - |
| | P63 | 22 | 21 | 0.881 | 41 | 0.3836 (ns) | 119.4 | 125.8 | 6.353 ± 7.213 | -8.215 to 20.92 | 0.269 (small) | - |
| | P112 | 23 | 26 | 3.058 | 10 | 0.0121 (*) | 107.5 | 77.22 | -30.27 ± 9.900 | -52.33 to -8.214 | 0.875 (large) | 28.16 |
| Gad1+ (GABAergic) [Fig. 6E-H] | P30 | 22 | 20 | 0.105 | 40 | 0.9172 (ns) | 167.0 | 168.2 | 1.195 ± 11.42 | -21.89 to 24.28 | 0.032 (trivial) | - |
| | P63 | 22 | 21 | 1.604 | 10 | 0.1398 (ns) | 144.9 | 156.8 | 11.96 ± 7.454 | -4.652 to 28.57 | 0.489 (small) | - |
| | P112 | 23 | 26 | 3.399 | 47 | 0.0014 (**) | 139.2 | 106.2 | -33.02 ± 9.715 | -52.56 to -13.48 | 0.973 (large) | 23.72 |
| Gad2+ (GABAergic) [Fig. 6I-L] | P30 | 22 | 20 | 0.414 | 10 | 0.6879 (ns) | 114.4 | 110.2 | -4.157 ± 10.05 | -26.55 to 18.24 | 0.128 (trivial) | - |
| | P63 | 22 | 21 | 1.273 | 41 | 0.2101 (ns) | 155.7 | 141.4 | -14.35 ± 11.27 | -37.10 to 8.411 | 0.388 (small) | - |
| | P112 | 23 | 26 | 0.995 | 10 | 0.3431 (ns) | 154.5 | 169.8 | 15.35 ± 15.44 | -19.03 to 49.75 | 0.285 (small) | - |
| GlyT2+ and/or Gad1+ and/or Gad2+ (Inhibitory) [Supp. Fig. 6] | P30 | 22 | 20 | 0.391 | 40 | 0.6975 (ns) | 246.7 | 252.7 | 5.968 ± 15.25 | -24.85 to 36.78 | 0.121 (trivial) | - |
| | P63 | 22 | 21 | 0.303 | 41 | 0.7634 (ns) | 260.3 | 263.5 | 3.158 ± 10.42 | -17.89 to 24.20 | 0.092 (trivial) | - |
| | P112 | 23 | 26 | 0.324 | 10 | 0.7529 (ns) | 217.2 | 212.5 | -4.694 ± 14.51 | -37.02 to 27.63 | 0.093 (trivial) | - |

**Table S3. Statistical details for assessment of excitatory populations.**

| Marker/s (Population) [Figure] | Time-point | n sections (N = 3 mice, 6 hemicords) | | Nested unpaired two-tailed t-test | | | wt mean | SOD1 mean | Descriptive report of differences | | Effect size (d) | Est. % loss |
|---|---|---|---|---|---|---|---|---|---|---|---|---|
| | | wt | SOD1 | t | df | P value | | | (SOD1-wt) ±SEM | 95% CI of diff. | | |
| Excitatory/Chx10$^+$ (V2a) [Fig. 7A-D] | P30 | 22 | 20 | 1.392 | 10 | 0.1941 (ns) | 19.11 | 16.05 | -3.059 ± 2.197 | -7.955 to 1.837 | 0.430 (small) | - |
| | P63 | 22 | 21 | 0.110 | 10 | 0.9150 (ns) | 15.70 | 15.96 | 0.2629 ± 2.400 | -5.085 to 5.611 | 0.034 (trivial) | - |
| | P112 | 23 | 26 | 4.894 | 47 | <0.0001 (****) | 21.74 | 9.423 | -12.32 ± 2.517 | -17.38 to -7.253 | 1.401 (large) | 57.57 |
| Excitatory/Shox2$^+$ (Shox2) [Fig. 7E-G] | P112 | 23 | 26 | 2.984 | 10 | 0.0137 (*) | 31.12 | 20.47 | -10.65 ± 3.569 | -18.60 to -2.699 | 0.854 (large) | 34.22 |
| Exc./Shox2$^+$/Chx10$^-$ (Shox2 non-V2a) [Fig. 7H] | P112 | 23 | 26 | 0.137 | 47 | 0.8914 (ns) | 13.96 | 13.65 | -0.3027 ± 2.205 | -4.739 to 4.133 | 0.039 (trivial) | - |
| Exc./Shox2$^+$/Chx10$^+$ (Shox2 V2a) [Fig. 7I-J] | P112 | 23 | 26 | 4.410 | 10 | 0.0013 (**) | 16.97 | 6.857 | -10.12 ± 2.294 | -15.23 to -5.005 | 1.262 (large) | 59.63 |
| Pitx2$^+$ ($V0_{C/G}$) [Fig. 8] | P30 | 22 | 20 | 0.214 | 40 | 0.8316 (ns) | 1.318 | 1.250 | -0.06818 ± 0.3186 | -0.712 to 0.576 | 0.066 (trivial) | - |
| | P63 | 22 | 21 | 0.738 | 10 | 0.4776 (ns) | 0.876 | 0.649 | -0.2266 ± 0.3071 | -0.911 to 0.458 | 0.225 (small) | - |
| | P112 | 23 | 26 | 0.271 | 10 | 0.7916 (ns) | 1.090 | 1.189 | 0.9940 ± 0.3663 | -0.717 to 0.916 | 0.078 (trivial) | - |
| Pitx2$^+$/ChAT$^+$ ($V0_C$) [Supp.Fig.8A,C,E,G] | P30 | 22 | 20 | 0.257 | 40 | 0.7989 (ns) | 0.864 | 0.800 | -0.06364 ± 0.2481 | -0.565 to 0.438 | 0.079 (trivial) | - |
| | P63 | 22 | 21 | 0.510 | 10 | 0.6215 (ns) | 0.736 | 0.593 | -0.1431 ± 0.2809 | -0.769 to 0.483 | 0.156 (trivial) | - |
| | P112 | 23 | 26 | 0.456 | 10 | 0.6579 (ns) | 0.907 | 0.771 | -0.1363 ± 0.2987 | -0.802 to 0.529 | 0.131 (trivial) | - |
| Pitx2$^+$/Vglut2$^+$ ($V0_G$) [Supp.Fig. 8B,D,F,H] | P30 | 22 | 20 | 1.765 | 40 | 0.0852 (ns) | 0.727 | 0.300 | -0.4273 ± 0.2421 | -0.917 to 0.062 | 0.545 (med.) | - |
| | P63 | 22 | 21 | 0.578 | 10 | 0.5762 (ns) | 0.333 | 0.208 | -0.1248 ± 0.2160 | -0.606 to 0.3565 | 0.176 (trivial) | - |
| | P112 | 23 | 26 | 0.294 | 10 | 0.7746 (ns) | 0.570 | 0.500 | -0.07018 ± 0.2385 | -0.602 to 0.4613 | 0.084 (trivial) | - |
| Vglut2$^+$ (Excitatory) [Fig. 9] | P30 | 22 | 20 | 1.557 | 40 | 0.1273 (ns) | 205.0 | 184.6 | -20.50 ± 13.16 | -47.10 to 6.106 | 0.481 (small) | - |
| | P63 | 22 | 21 | 1.832 | 30 | 0.0742 (ns) | 149.7 | 177.9 | 28.20 ± 15.39 | -67.12 to 10.72 | 0.559 (med.) | - |
| | P112 | 23 | 26 | 0.099 | 10 | 0.9225 (ns) | 188.0 | 190.2 | 2.130 ± 21.37 | -45.48 to 49.74 | 0.028 (trivial) | - |
| Ventral ChAT$^+$ (Motor neurons) [Fig. 10] | P30 | 22 | 20 | 0.205 | 10 | 0.8417 (ns) | 20.59 | 19.95 | -0.6323 ± 3.085 | -7.505 to 6.241 | 0.063 (trivial) | - |
| | P63 | 22 | 21 | 0.929 | 10 | 0.3746 (ns) | 15.86 | 13.55 | -2.312 ± 2.488 | -7.855 to 3.232 | 0.283 (small) | - |
| | P112 | 23 | 26 | 2.348 | 47 | 0.0231 (*) | 9.826 | 7.038 | -2.788 ± 1.187 | -5.176 to -0.3996 | 0.672 (med.) | 28.37 |
